# Practical parameter identifiability for spatiotemporal models of cell invasion

**DOI:** 10.1101/839282

**Authors:** Matthew J Simpson, Ruth E Baker, Sean T Vittadello, Oliver J Maclaren

## Abstract

We examine the practical identifiability of parameters in a spatiotemporal reaction-diffusion model of a scratch assay. Experimental data involves fluorescent cell cycle labels, providing spatial information about cell position and temporal information about the cell cycle phase. Cell cycle labelling is incorporated into the reaction–diffusion model by treating the total population as two interacting subpopulations. Practical identifiability is examined using a Bayesian Markov chain Monte Carlo (MCMC) framework, confirming that the parameters are identifiable when we assume the diffusivities of the subpopulations are identical, but that the parameters are practically non-identifiable when we allow the diffusivities to be distinct. We also assess practical identifiability using a profile likelihood approach, providing similar results to MCMC with the advantage of being an order of magnitude faster to compute. Therefore, we suggest that the profile likelihood ought to be adopted as a screening tool to assess practical identifiability before MCMC computations are performed.

## 1. Introduction

Combined cell migration and cell proliferation leads to moving fronts of cells, often called *cell invasion* [1], which is essential for tissue repair and wound healing [2]. While sophisticated experimental techniques are continually developed to interrogate cell invasion, traditional *scratch assays* remain widely used because they are simple, fast and inexpensive [3]. Scratch assays involve growing (Figure 1a-b) and then scratching (Figure 1c) a cell monolayer, and imaging a small region around the scratch as the wounded area closes (Figure 1d-e).

**Figure 1:**
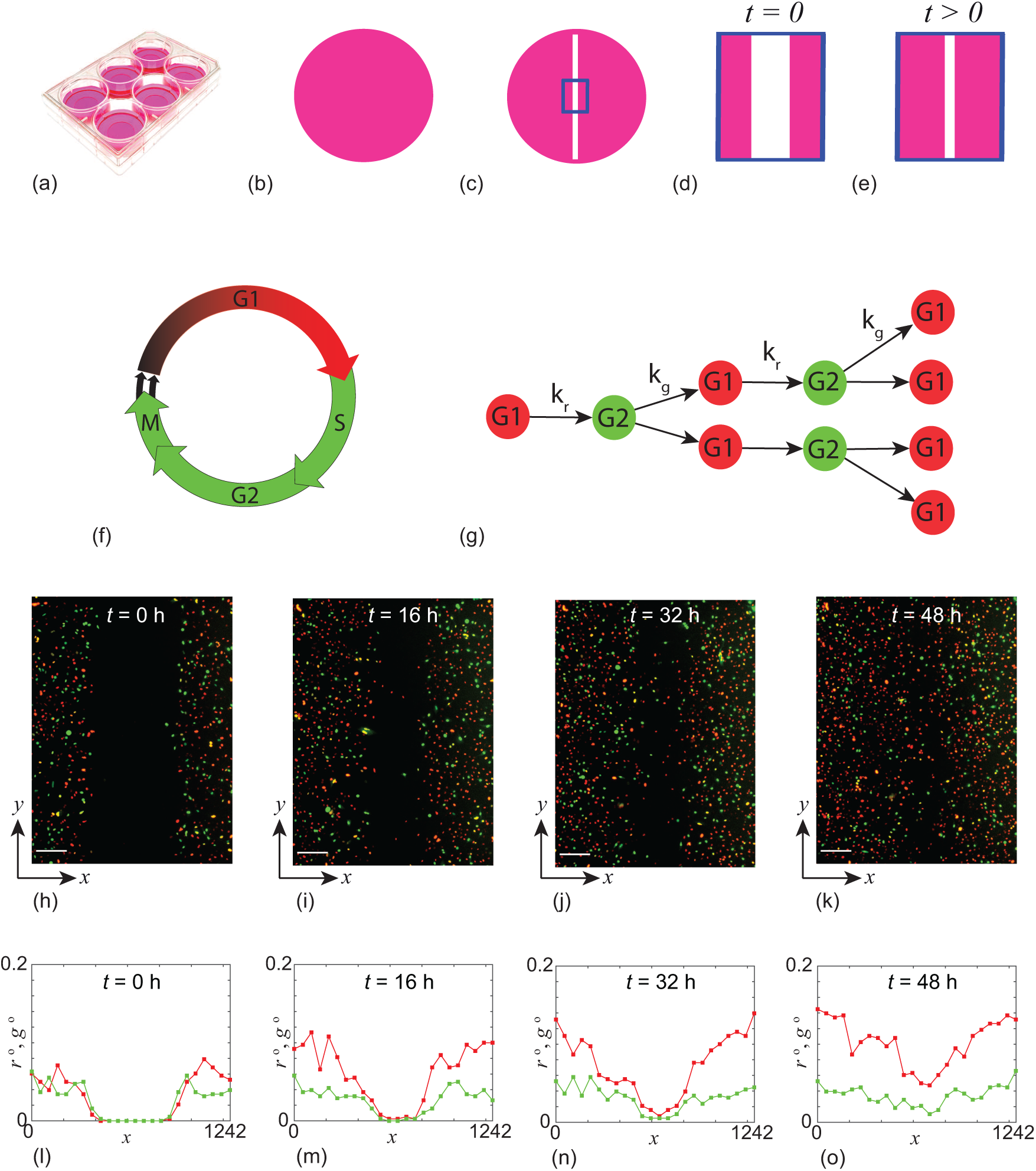
Experiments in a 6-well plate (a) are initiated with a uniform cell monolayer (b). A scratch is made (c), and the 1296 *µm* × 1745 *µ*m field of view is observed between *t* = 0 h (d) and *t* = 48 h (e). The cell cycle is labelled using fluorescent ubiquitination-based cell cycle indicator (FUCCI) [29] so that cells in G1 phase fluoresce red and cells in S/G2/M phase fluoresce green (f). A freely-cycling cell in G1 phase (red) will transition into S/G2/M phase (green) at rate *k*_*r*_ > 0. A freely-cycling cell in S/G2/M phase (green) will divide into two cells in G1 phase (red) at rate *k*_*g*_ > 0, provided there is sufficient space to accommodate division (g). Images of the field–of–view at *t* = 0, 16, 32 and 48 h (h–k), with the nondimensional densities of G1 (red) and S/G2/M (green) subpopulations shown as a function of space and time (l–o). Scale bars in (l–o) show 200 *µ*m.

The first mathematical models to interpret experimental observations of cell invasion were reaction–diffusion models [4]. Sherratt and Murray [5] modelled a wound-healing-type assay using the Fisher-Kolmogorov model, and many subsequent studies have related solutions of mathematical models to experimental observations, including Maini et al. [2], Sengers et al., [6] and Nardini et al. [7]. By matching the solution of a mathematical model with certain experimental observations, these studies provide both parameter estimates and mechanistic insight.

One challenge of using scratch assays is that there is no standard, widely accepted experimental protocol. There are many differences in: (i) the initial monolayer density [8]; (ii) the width and shape of the scratch [1, 9]; and, (iii) the experimental timescale [3]. These differences make it is difficult to compare different experiments, and it is unclear whether different mathematical models of varying complexity (i.e. varying numbers of unknown parameters or varying the model structure) are required under different experimental protocols.

It is reasonable to argue that simple experimental data ought to be modelled using simple mathematical models with a small number of unknown parameters, and that more sophisticated mathematical models with additional parameters should be used only when greater amounts of experimental data are available. However, it is difficult to use this kind of qualitative reasoning to justify particular decisions about how complicated a mathematical model ought to be. Methods of *identifiability analysis* [10–19, 22] provide more systematic means to determine the appropriate model complexity relative to available data. A model is *identifiable* when distinct parameter values imply distinct distributions of observations, and hence when it is possible to uniquely determine the model parameters using an infinite amount of ideal data [23–26]. In the systems biology literature that focuses on models formulated as ordinary differential equations, *identifiability* has also been referred to as *structural identifiability* [18, 19]. Working with ordinary differential equations that describe temporal processes, it is possible to formally analyse the structural identifiability [18–21]. In contrast, here we work with reaction–diffusion partial differential equations since we are interested in spatiotemporal processes. In this case, formal analysis of structural identifiability is more difficult and so we explore structural identifiability numerically (Supplementary Material).

In contrast to predictive performance measures, like Akaike’s Information Criterion (AIC) [27], identifiability analysis focuses on distinguishing parameter values and, as a result, understanding underlying mechanisms. Shmueli characterises this general difference as the choice of whether to ‘explain or predict’ [28]. The formal definition of identifiability requires an infinite amount of ideal data. Thus, in addition to the strict technical concept of (structural) identifiability, the terms practical *identifiability* [15] and *estimability* [23, 26] are used to describe whether it is possible to provide reasonably precise estimates using finite, non-ideal data.

Since practical identifiability is a finite, non-ideal data concept, it depends more strongly on the inferential framework adopted and is less clearly defined in the literature than structural identifiability. We consider practical identifiability analysis to encompass a set of *precision-based* pre-data planning and post-data analysis tools similar to those advocated by Cox [30], Bland [31] and Rothman and Greenland [32]. This is consistent with the systems biology literature [10–17, 22], where it is typically characterised in terms of the ability to provide finite, sufficiently precise interval estimates at various levels of confidence or credibility given finite amounts of non-ideal, real data. This is the approach we take here; we consider both the typical precision achieved under repeated simulation of ideal synthetic data (Supplementary Information) and the post-data, realised precision given real data (Main Document).

In systems biology, experimental data often take the form of time series describing temporal variations of molecular species in a chemical reaction network and these data are modelled using ordinary or stochastic differential equations [33]. Here, we focus on applying methods of practical identifiability analysis to spatiotemporal partial differential equation models, which represent both spatial and temporal variations in quantities of interest. It is instructive to consider both a Bayesian inference framework [13, 14] and profile likelihood analysis [15, 16].

Bayesian inference is widely adopted within the mathematical biology community [34–39]. A drawback of the Bayesian approach is that it can be computationally expensive, particularly when Markov chain Monte Carlo (MCMC) methods are used to sample the distributions of interest. In contrast, profile likelihood analysis is a more standard tool of statistical inference [30, 40], but is far less familiar within the mathematical biology community. Being optimisation-based, profile likelihood can be computationally inexpensive.

Here we consider data from a scratch assay where cells are labelled using two fluorescent probes to show realtime progression through the cell cycle [29]. Such labelling allows us to describe the total population of cells as two subpopulations according to the two cell cycle labels [41]. This experimental approach provides far more information than a standard scratch assay without cell cycle labels. We model the experiments using a system of reaction–diffusion equations that are an extension of the Fisher–Kolmogorov model. Using this data and this model, we explore the practical identifiability of the model parameters using both Bayesian inference and profile likelihood.

One of the main outcomes of this work is that, instead of performing potentially computationally expensive Bayesian inference to assess practical identifiability and determine parameter estimates, we show that a profile likelihood analysis can provide a fast and reliable preliminary assessment of practical identifiability. Both analysis methods provide good insight into the identifiability of the spatiotemporal models considered here. In summary, we consider a minimal model scratch assay data generated using fluorescent cell cycle labelling and show that it is practically identifiable, whereas a simple extension of that model is not, despite the fact that numerical experiments indicate that it is structurally identifiable in the limit of infinite data. In all cases, the Bayesian analysis is consistent with profile likelihood results. However, the profile likelihood analysis is an order of magnitude faster to implement.

## 2. Experimental data

We consider a scratch assay performed with the 1205Lu melanoma cell line with FUCCI labelling [29]. The scratch assay is initiated by seeding 1205Lu melanoma cells into 6-well plates and allowing sufficient time for the cells to attach, begin to proliferate (Figure 1a), and to form an approximately uniform monolayer (Figure 1b). A scratch is made in the monolayer (Figure 1c) and a small region of the experiment is imaged (Figure 1c-d) [41]. The geometry of the experiment means that the distribution of cells in the imaged region is independent of vertical position, and so we treat the cell density as a function of horizontal position, *x*, and time, *t* [41]. The boundaries around the imaged region are not physical boundaries and cells are free to migrate across these boundaries. Since the density of cells is approximately constant away from the scratch, the net flux of cells locally across each boundary of the imaged region is approximately zero [41].

Cells in G1 phase fluoresce red and cells in S/G2/M phase fluoresce green (Figure 1f). We represent FUCCI labelling by treating the total population as two subpopulations: (i) red cells in G1, and (ii) green cells in S/G2/M. Cells in G1 (red) transition into S/G2/M cells (green) at rate *k*_*r*_ > 0. This red-to-green transition does not involve any change in cell number so we assume this transition is unaffected by the availability of physical space. Cells in S/G2/M (green) undergo mitosis at rate *k*_*g*_ > 0 to produce two daughter cells in G1 (red) (Figure 1f). Since the green-to-red transition involves the production of two daughter cells we assume this transition only occurs provided there is sufficient space.

Images of the experiment (Figure 1h-k) show that cells move into the initially–scratched region while simultaneously progressing through the cell cycle. At the beginning of the experiment the scratch is approximately 500 *µ*m wide, and by the end of the experiment the initially-vacant wound space becomes occupied as cells have migrated into the gap. Visual examination of the density of cells away from the scratched region shows that the cell density increases with time, and this is driven by cell proliferation. To quantify these visual observations we note that each experimental image is 1296 *µ*m wide, and we discretise each image into 24 equally–spaced columns of width 54 *µ*m. We count the number of red cells and green cells per column, divide by the area of the column and by the theoretical maximum packing density of 0.004 cells/*µ*m^2^ [41] to give an estimate of the nondimensional density of red cells and green cells per column. Assuming that each experimental density estimate represents the centre of each column, we plot these data (Figure 1l–o) shows the spatial variation in density of G1 (red) and S/G2/M (green) cells as a function of space and time.

We summarise the experimental data using the following notation. Our observations are vectors containing the observed nondimensional density of red cells, 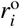, and the observed nondimensional density of green cells, 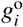, per column at each position *x*_*i*_ and time *t*_*i*_, i.e.

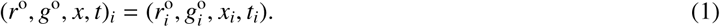

We use an explicit superscript ‘o’ to distinguish (noisy) observations of *r* and *g* from their modelled counterparts introduced below, while we assume that the space and time coordinates *x* and *t*, respectively, are noise-free. In particular we use

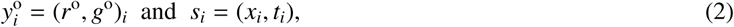

to denote the *i*th combined vector of observations of *r* and *g*, and the *i*th combined vector of spatiotemporal coordinates *x* and *t*, respectively. Observations are taken at 24 equally–spaced spatial locations, where the spacing between the observations is 54 *µ*m, at four equally–spaced time points, *t* = 0, 16, 32 and 4 h. Thus *y*^o^ consists of 24 × 4 = 96 observations of both of *r* and *g*, giving *N* = 24 × 4 × 2 = 192 measurements of cell density in total. Experimental data are provided (Supplementary Material).

## 3. Mathematical model

We treat our *mathematical model* as having two components: (i) a deterministic spatiotemporal *process* model governing the dynamics of the experiment; and, (ii) a probabilistic *observation* model. This approach invokes the reasonable assumption that observations are noisy versions of a deterministic latent model [13]. We describe both models and note that the same model components are used in the Bayesian and the profile likelihood analyses.

### 3.1. Process model

A reasonably simple process model for the FUCCI scratch assay can be written as

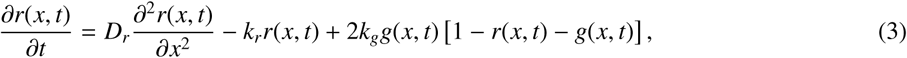

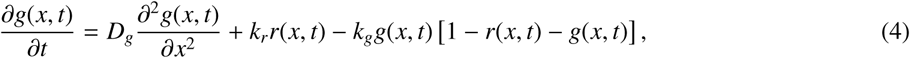

where *D*_*r*_ > 0 is the diffusivity of cells in G1, *D*_*g*_ > 0 is the diffusivity of cells in S/G2/M, *k*_*r*_ > 0 is the rate at which cells in G1 transition into S/G2/M, and *k*_*g*_ > 0 is the rate at which cells in S/G2/M undergo mitosis to produce two cells in G1 [41]. The solution of Equations (3)–(4), *r*(*x, t*) and *g*(*x, t*), are taken to represent the (continuous) underlying mean densities, while 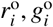 represent noisy observations of these mean densities at the measurement locations *s*_*i*_ = (*x*_*i*_, *t*_*i*_).

The process model has four parameters: *θ* = (*D*_*r*_, *D*_*g*_, *k*_*r*_, *k*_*g*_). Given suitable initial conditions, *r*(*x*, 0) and *g*(*x*, 0), and parameter values *θ*, Equations (3)–(4) are solved numerically (Supplementary Material).

Despite the abundance of experimental data in Figure 1 and the apparent simplicity of Equations (3)–(4), it is unclear whether these experimental data are sufficient estimate the four parameters: *θ* = (*D*_*r*_, *D*_*g*_, *k*_*r*_, *k*_*g*_). Therefore, we also consider a simpler model whereby we set *D* = *D*_*r*_ = *D*_*g*_, implying that cells in G1 diffuse at the same rate as cells in S/G2/M. This simpler model is characterised by three parameters: *θ* = (*D, k*_*r*_, *k*_*g*_).

### 3.2. Observation model

We assume that observations 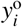 are noisy versions of the latent model solutions, *y*(*s*_*i*_, *θ*). A standard approach in the mathematical biology literature is to further assume that this observation error is independent and identically–distributed (iid), and that this noise is additive and normally–distributed, with zero mean and common variance *σ*^2^ [8, 13]. Therefore, we have

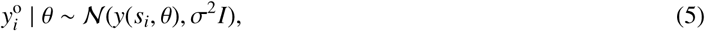

where *I* is the 2×2 identity matrix. Since our main interest is in *θ* = (*D*_*r*_, *D*_*g*_, *k*_*r*_, *k*_*g*_), we directly specify *σ* = 0.05 [8, 13], and we will discuss this choice later.

## 4. Practical identifiability analysis

We consider two approaches for practical identifiability analysis. The first is based on Bayesian inference and MCMC, and follows the approach outlined by Hines et al. [13]. The second approach is based on profile likelihood and follows Raue et al. [15]. Both approaches have been considered in the context of identifiability of non-spatial models, and, as we will show, the same ideas can deal with models of spatiotemporal processes.

As discussed by Raue et al. [42], estimation results from MCMC and profile likelihood are typically similar in the case of identifiable parameters (and in the low-dimensional setting), but can be very different in the presence of non-identifiability. In particular, Raue et al. [42] demonstrate that results from MCMC can be potentially misleading in the presence of non-identifiability. On the other hand, Hines et al. [13] and Seikmann et al. [14] argue that appropriate MCMC diagnostics can indicate the presence of non-identifiability. Hence we use both methods and consider the extent to which they agree.

### 4.1. Parameter bounds

Before outlining the identifiability methodologies, we note that the parameters in Equations (3)–(4) have a physical interpretation and we can formulate some biologically–motivated bounds. Practical identifiability, given observed data, can be evaluated by comparing the width of (realised) interval estimates relative to these simple bounds.

Previous estimates of melanoma cell diffusivity in similar experiments found that typical values are often less than 1000 *µ*m^2^ /h [43] so we take conservative bounds on *D*_*r*_ and *D*_*g*_ to be 0 < *D*_*r*_, *D*_*g*_ < 2000 *µ*m^2^ /h. Bounds on *k*_*r*_ and *k*_*g*_ can be inferred from previous experimental measurements. The duration of time 1205Lu melanoma cells remain in the G1 phase varies between 8–30 h, whereas the time 1205Lu melanoma cells remain in the S/G2/M phase varies between 8–17 h [29]. These measurements imply 0.033 < *k*_*r*_ < 0.125 /h and 0.059 < *k*_*g*_ < 0.125 /h, so we take conservative bounds on *k*_*r*_ and *k*_*g*_ to be 0 < *k*_*r*_, *k*_*g*_ < 0.2 /h.

### 4.2. Bayesian inference

Following Hines et al. [13] we take a Bayesian approach to assess parameter identifiability. Bayesian inference relies on Bayes’ theorem, written here as

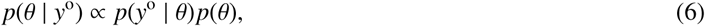

where *p*(*y*^o^ | *θ*) is the *likelihood function, p*(*θ*) is the *prior* and *p*(*θ* | *y*^o^) is the *posterior*. The posterior distribution is the inferential target; this summarises the information about the parameter *θ* in light of the observed data, *y*^o^, and the prior information specified by *p*(*θ*). The likelihood function represents the information contributed by the data, and corresponds to the probabilistic observation model in 5 evaluated at the observed data.

We explore the parameter space *θ* by sampling its posterior distribution using a Metropolis-Hastings MCMC algorithm [13]. The Markov chain starts at position *θ*_*i*_, and a potential move to *θ*^*^ is accepted with probability *α*,

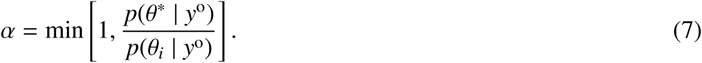

This Markov chain tends to move toward regions of high posterior probability, but it also provides a mechanism to move away from local minima by allowing transitions to regions of lower posterior probability. The Markov chain produced by this algorithm explores the parameter space in proportion to the posterior probability and provides a finite number of independent, identically distributed samples from the posterior distribution. Proposals in the MCMC algorithm are made by sampling a multivariate normal distribution with zero mean and covariance matrix Σ. When *θ* = (*D*_*r*_, *D*_*g*_, *k*_*r*_, *k*_*g*_) we specify Σ = diag(10^2^, 10^2^, 10^−6^, 10^−6^), whereas when *θ* = (*D, k*_*r*_, *k*_*g*_) we specify Σ = diag(10^2^, 10^−6^, 10^−6^).

Poor identifiability in a Bayesian setting using MCMC is typically characterised by poorly converging chains, label-switching, multimodal or overly-broad distributions, and similar phenomena that can be diagnosed either graphically or by computing various diagnostic statistics [13, 14]. We discuss these diagnostic methods in more detail in the Results section.

### 4.3. Profile likelihood

Here we describe how a profile likelihood identifiability analysis can be undertaken. This is based on the same likelihood as above, *p*(*y*^o^ | *θ*), but here we will present results in terms of the normalised likelihood function, denoted

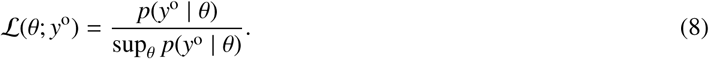

This is considered as a function of *θ* for fixed data *y*^o^. It is also common to take the likelihood as only defined up to a constant, but here we fix the proportionality constant by presenting results in terms of the normalised likelihood function as above.

We assume our full parameter *θ* can be partitioned into an *interest* parameter *ψ* and *nuisance* parameter *λ*, i.e. *θ* = (*ψ, λ*). These can also be considered functions *ψ*(*θ*) and *λ*(*θ*) of the full parameter *θ*, specifying the partition components. Here we will restrict attention to a scalar interest parameter and a vector nuisance parameter, i.e. multiple scalar nuisance parameters. Then the profile likelihood for the interest parameter *ψ* can be written as [30, 40]

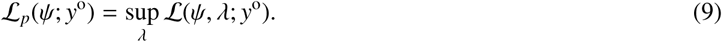

In Equation (9), *λ* is ‘optimised out’ for each value of *ψ*, and this implicitly defines a function *λ*^*^(*ψ*) of optimal *λ* values for each value of *ψ*. For example, given the full parameter *θ* = (*D*_*r*_, *D*_*g*_, *k*_*r*_, *k*_*g*_), we may consider the diffusion constant of red cells as the interest parameter and the other parameters as nuisance parameters, i.e. *ψ*(*θ*) = *D*_*r*_ and *λ*(*θ*) = (*D*_*g*_, *k*_*r*_, *k*_*g*_). This would give

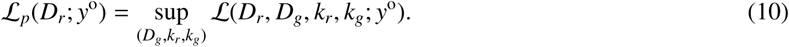

Given the assumption of normally distributed errors, and assuming a fixed, common standard deviation *σ*, profiling reduces to solving a series of nonlinear least squares problems, one for each value of the parameter of interest, by using log-density functions. Additional details and example calculations are provided in the Supplementary Material.

We implement this optimisation using MATLAB’s *lsqnonlin* function with the ‘trust-region-reflective’ algorithm to implement bound constraints [44]. For each value of the interest parameter, taken over a sufficiently fine grid, the nuisance parameter is optimised out and the previous optimal value is used as the starting guess for the next optimisation problem. Uniformly–spaced grids of 50 points, defined between previously–discussed lower and upper bounds are used. Results are plotted in terms of the normalised profile likelihood functions; cubic interpolations are used to define the profiles for all parameter values. Solutions did not exhibit any notable dependence on initial guesses.

The likelihood function is often characterised as representing the information that the data contains about the parameters, and the relative likelihood for different parameter values as indicating the relative evidence for these parameter values [40, 45, 46]. As such, a flat profile is indicative of non-identifiability, therefore a lack of information in the data about a parameter [15]. In general, the degree of curvature is related to the inferential precision [15, 16, 42]. Likelihood-based confidence intervals can be formed by choosing a threshold-relative profile likelihood value, which can be approximately calibrated via the chi-square distribution (or via simulation). We use a threshold of 0.15 as a reference, which corresponds to an approximate 95% confidence interval for sufficiently regular problems [40]. The points of intersection were determined using the interpolated profile likelihood functions.

## 5. Results and Discussion

Here we consider the results of the above methods of identifiability analysis under the two scenarios introduced previously. In Scenario 1 we consider Equations (3)–(4) under the assumption that both subpopulations have the same diffusivity, *D*_*r*_ = *D*_*g*_ = *D*, so that *θ* = (*D, k*_*r*_, *k*_*g*_). In Scenario 2 we consider the same model without this assumption so that *θ* = (*D*_*r*_, *D*_*g*_, *k*_*r*_, *k*_*g*_). We first consider the Bayesian analysis using MCMC, and then the profile likelihood analysis. All units of diffusivities are *µ*m^2^/h and all units for rate parameters are /h.

### 5.1. Bayesian analysis

As we are focused on the question of identifiability, here we use uniform priors on all parameters. This is a natural choice when we are interested in identifiability since we want to focus on the information about the parameter values inherent within the data rather than from some imposed prior [13].

In the first scenario where *D*_*r*_ = *D*_*g*_ = *D*, we see rapid convergence of the Markov chain for *D, k*_*r*_ and *k*_*g*_ (Figure 2a-c). Importantly, after the Markov chain moves away from the initial location, *θ*_0_, it remains within the previously–stated conservative bounds. Additional results in the Supplementary Material confirm similar results for different *θ*_0_. A plot matrix representation of the univariate and bivariate marginal densities is given (Figure 2d-i) [47] where the univariate marginals (Figure 2d, g, i) are unimodal and approximately symmetric. The univariate posterior modes are 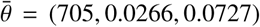, confirming that each component lies within the biologically–feasible range, and the univariate 95% credible intervals are relatively narrow. A posterior predictive check [47] with 30 randomly sampled parameter choices from the converged region of the Markov chain provides visual confidence in the MCMC results and the appropriateness of the model (Figure 2j-m) [47]. Additional comparison of the experimental data and solution of the model parameterised with 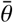 also confirms this (Supplementary Material). In summary, setting *D* = *D*_*r*_ = *D*_*g*_, our model parameters appear to be identifiable according to the Bayesian approach.

**Figure 2:**
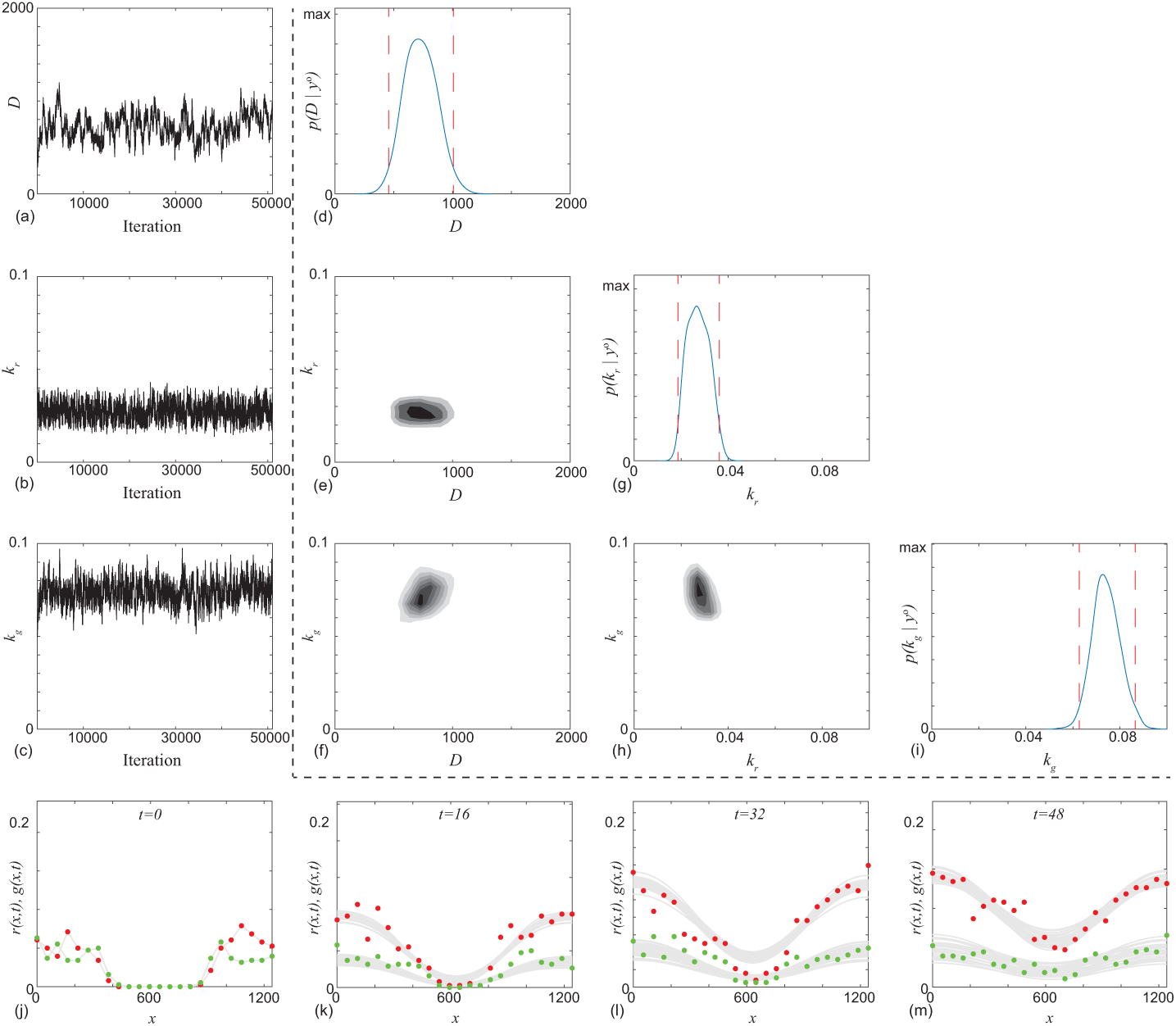
Typical Markov chain iterations, of length 51,000, for *D, k*_*r*_ and *k*_*g*_ in (a)–(c), respectively. In this case the Markov chain is initiated with *θ*_0_ = (500, 0.05, 0.05). Results in (d)–(i) show a plot matrix representation of the univariate marginals and bivariate marginals estimated using the final 50,000 iterations of the Markov chain in (a)–(c). For the univariate distribution the posterior modes are 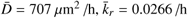, and 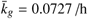, and the 95% credible intervals are *D* ∈ [455, 1007], *k*_*r*_ ∈ [0.0187, 0.0362] and *k*_*g*_ ∈ [0.0628, 0.0727]. In the univariate marginals the 95% credible intervals are shown in red dashed vertical lines, in the bivariate marginals the region of maximum density is shown in the darkest shade. Results in (j)–(m) superimpose 30 solutions of Equations (3)–(4) where *θ* is randomly sampled from the Markov chain. MCMC results use *σ* = 0.05.

In the second scenario, where *D*_*r*_ ≠ *D*_*g*_, the Markov chains for *D*_*r*_, *D*_*g*_, *k*_*r*_ and *k*_*g*_ (Figure 3a-d) are very different to the first scenario. The Markov chains for *k*_*r*_ and *k*_*g*_ remain within biologically–feasible bounds. However, the Markov chains for *D*_*r*_ and *D*_*g*_ often move beyond the previously-stated conservative bounds with no obvious convergence. Additional results for different choices of *θ*_0_ (Supplementary Material) indicate that these poor results for *D*_*r*_ and *D*_*g*_ are consistent across a number of choices of *θ*_0_. A plot matrix representation confirms that we obtain well–behaved posterior distributions for *k*_*r*_ and *k*_*g*_, but that the results for *D*_*r*_ and *D*_*g*_ are problematic since the univariate posterior distributions are not unimodal and the 95% credible intervals are relatively wide (Figure 3e–n). A posterior predictive check with 30 randomly sampled parameter choices from the latter part of the Markov chain are provided for completeness (Figure 3o–r). The posterior predictive check compares model predictions with the experimental data, with additional comparisons using the posterior median (Supplementary Material). However, these comparisons are less useful since they are based on parameters arising from a Markov chain that has not settled to a well-defined posterior. In summary, without assuming *D*_*r*_ = *D*_*g*_, our model parameters are practically non-identifiable according to the Bayesian approach.

**Figure 3:**
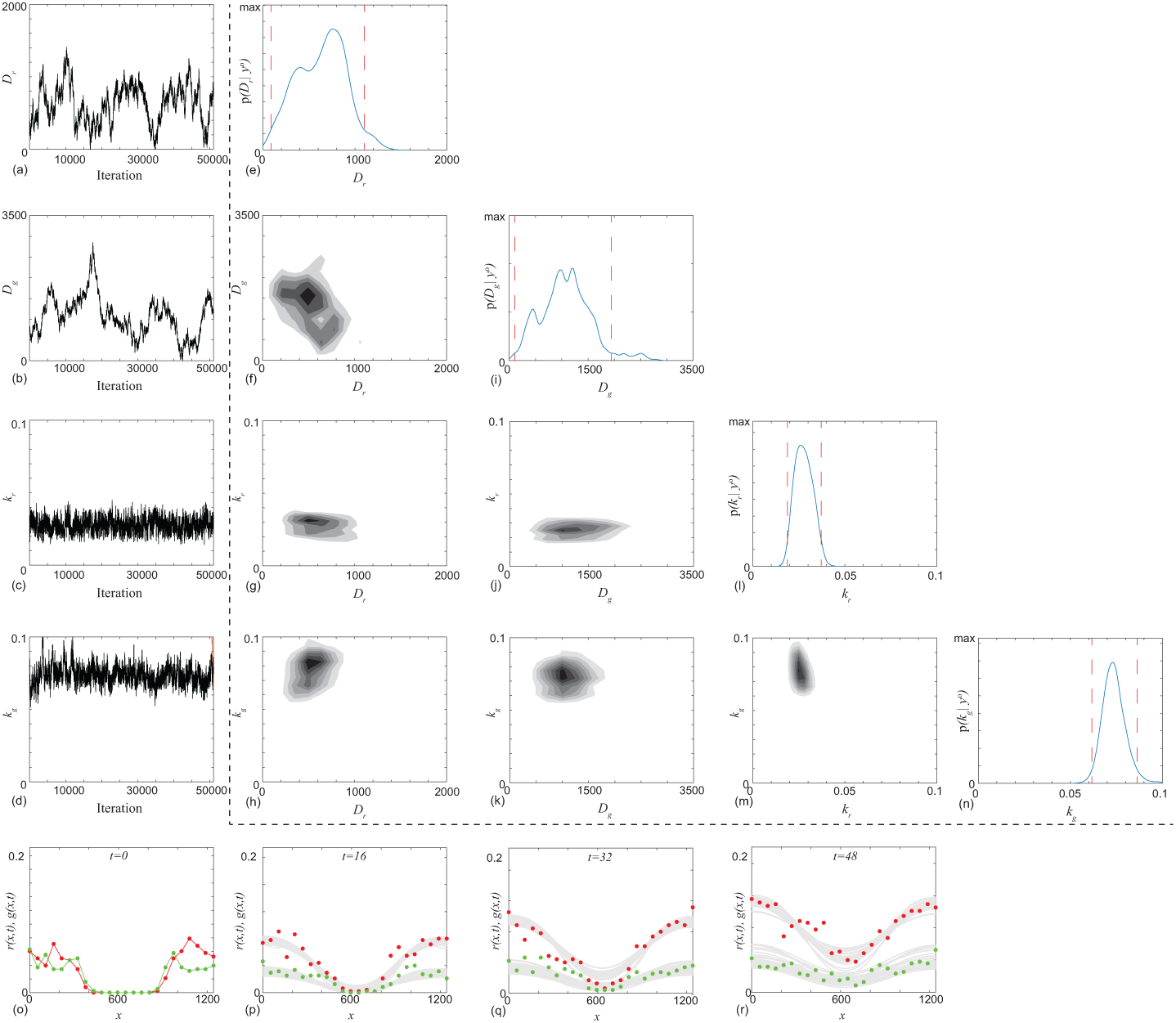
Typical Markov chain iterations, of length 51,000, for *D*_*r*_, *D*_*g*_, *k*_*r*_ and *k*_*g*_ in (a)–(d), respectively. In this case the Markov chain is initiated with *θ*_0_ = (228, 803, 0.0334, 0.0790). Results in (e)–(n) show a plot matrix representation of the univariate marginals and bivariate marginals estimated using the final 50,000 iterations of the Markov chain in (a)–(d). For the univariate distribution the posterior modes are 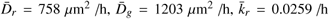, and 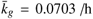. The 95% credible intervals are *D*_*r*_ ∈ [86, 1100], *D*_*g*_ ∈ [104, 1939], *k*_*r*_ ∈ [0.0188, 0.0371] and *k*_*g*_ ∈ [0.0617, 0.0863]. In the univariate marginals the 95% credible intervals are shown in red vertical dashed lines, in the bivariate marginals the region of maximum density is shown in the darkest shade. Results in (o)–(r) superimpose 30 solutions of Equations (3)–(4) where *θ* is randomly sampled from the Markov chain. MCMC results use *σ* = 0.05.

### 5.2. Profile likelihood analysis

Here we consider the results from our profile likelihood analysis. Each profile corresponds to taking a particular parameter as the interest parameter and the remaining parameters as nuisance parameters to be optimised out. More general partitions into an interest parameter and nuisance parameters are also considered in the Supplementary Material. Here we present results for Scenario 1 (Figure 4a–c) and results for Scenario 2 (Figure 4d–f).

**Figure 4:**
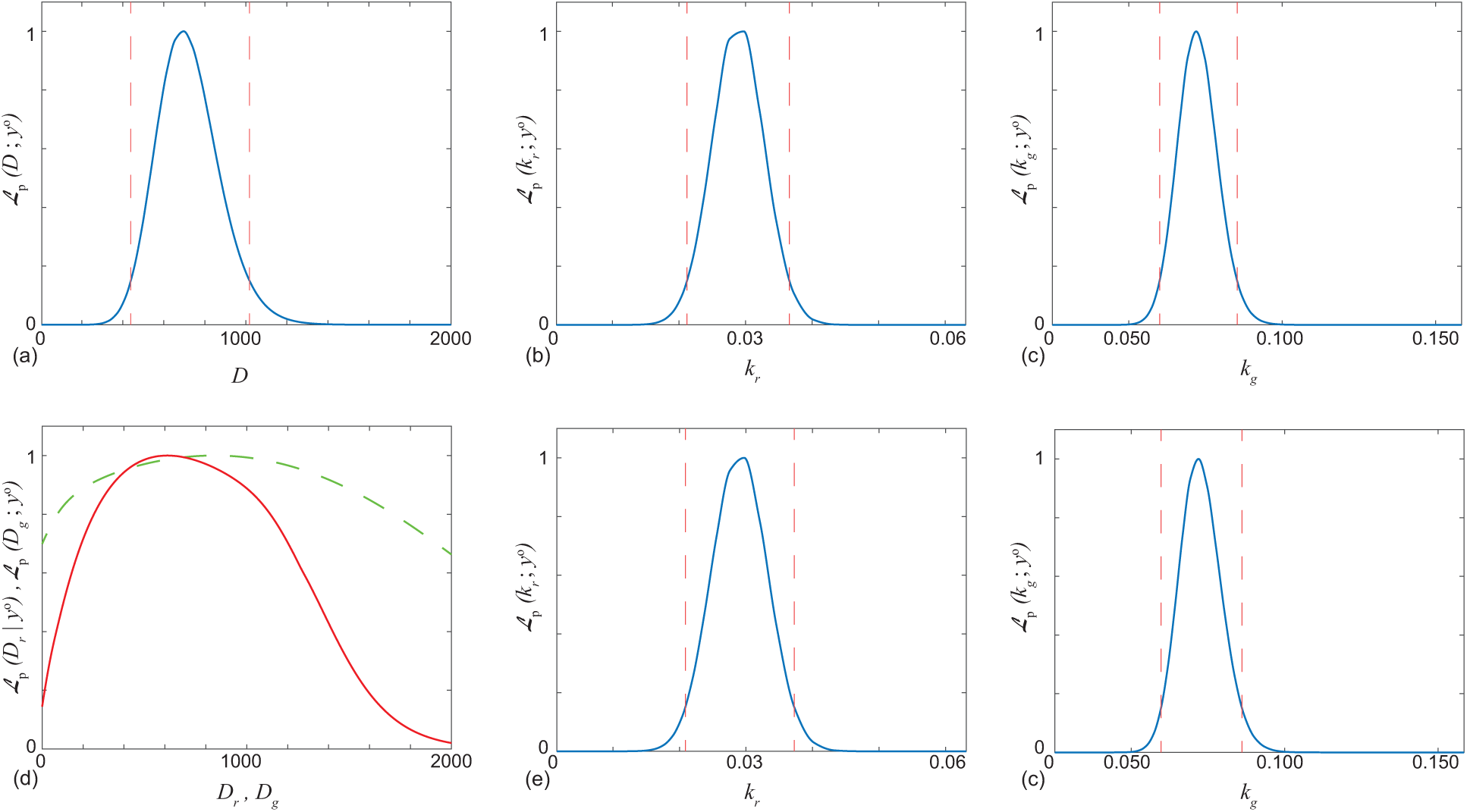
Profile likelihoods. Each profile likelihood is computed using a grid of 50 equally-spaced points; cubic interpolation iws used for display and to determine confidence intervals. Approximate 95% confidence intervals are indicated based on a relative likelihood threshold of 0.15 [40]. Results for the first scenario where *D*_*r*_ = *D*_*g*_ = *D* (a-c), and results for the second scenario where *D*_*r*_ ≠ *D*_*g*_ (d-f). In (d) the profile likelihood for *D*_*g*_ and *D*_*r*_ are in red (solid) and green (dashed), respectively. Interval estimates are: *D* ∈ [439, 1018], *k*_*r*_ ∈ [0.0212, 0.0366], *k*_*g*_ ∈ [0.0601, 0.0853] for *D* = *D*_*r*_ = *D*_*g*_. Maximum likelihood estimates are: 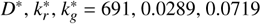 for *D* = *D*_*r*_ = *D*_*g*_. Interval estimates are: *D*_*r*_ ∈ [1.4, 1647], *D*_*g*_ ∈ [0, 2000] (i.e. the whole interval), *k*_*r*_ ∈ [0.0209, 0.0373], *k*_*g*_ ∈ [0.0597, 0.0861] for *D*_*r*_ ≠ *D*_*g*_. Maximum likelihood estimates are: 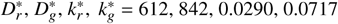 for *D*_*r*_ ≠ *D*_*g*_.

In the first scenario where *D*_*r*_ = *D*_*g*_ = *D*, we see regular shaped profiles with clearly defined peaks for *D, k*_*r*_ and *k*_*g*_ (Figure 4a-c). These results are very similar to those obtained from the Bayesian analysis using MCMC, both in terms of interval estimates and maximum likelihood estimates. The maximum likelihood estimates are 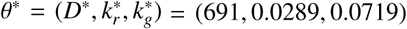.

In the second scenario, where *D*_*r*_ ≠ *D*_*g*_, we see very similar estimates for *k*_*r*_ and *k*_*g*_. The diffusivities, *D*_*r*_ and *D*_*g*_, are much less well-determined, however. In particular, the profile-likelihood-based confidence interval for *D*_*g*_ covers the entire range of the previously-stated conservative bounds, while that for *D*_*r*_ only excludes a negligible part of this region. Thus, while *k*_*r*_ and *k*_*g*_ are well identified, *D*_*r*_ and *D*_*g*_ are not. These results are again very similar to those obtained from the Bayesian analysis using MCMC, though the diffusivity estimates are even more conservative here. The maximum likelihood estimates are 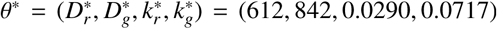. We also generated profiles for the difference *D*_*r*_ − *D*_*g*_ (Supplementary Material). The associated interval encompasses a broad range of values, including zero, but also includes very large positive and negative values. This is consistent with a lack of practical identifiability of *D*_*r*_ − *D*_*g*_. Despite the fact that the diffusivities are practically non-identifiable with our relatively abundant experimental data, further experimentation with synthetic data indicates that all parameters are, in fact, strictly identifiable provided sufficient data (Supplementary Material).

In both cases the results of the profile likelihood analysis is consistent with the MCMC-based Bayesian analysis. However, the profiles are far cheaper to compute than full MCMC posterior distributions. For example, each profile here was computed in less than one minute on a standard laptop whereas the MCMC analysis took approximately 30 minutes of computation. Each optimisation for a given value of the target parameter in the profile analysis took less than one second to solve, and profiles can be computed in parallel.

## 6. Conclusions and Outlook

In this work we assess the practical identifiability of a spatiotemporal reaction–diffusion model of a scratch assay where we have access to a relatively large amount of experimental data. In the literature, most scratch assays are reported by providing single images at the beginning and the conclusion of the experiment without making measurements of spatial distributions of cell density at multiple time points (e.g. Fattahi et al. [48]; Wang et al. [49]). In contrast, we consider detailed cell density measurements of two subpopulations at four time points with a relatively high spatial resolution of cell density at 24 spatial locations. This means that we work with 24 × 4 × 2 = 192 measurements of cell density. Given this data set, a key decision when using a mathematical model is to determine how complicated the model ought to be. We first explored this question through a Bayesian, MCMC-based framework, and then with a profile likelihood–based approach.

MCMC–based Bayesian analysis is relatively common in the systems biology literature for temporal models [34]. Bayesian MCMC approaches for spatiotemporal models are less common, but increasingly of interest. While it is established that practical parameter identifiability can often be diagnosed using an MCMC framework, many applications of Bayesian MCMC in the mathematical biology literature never explicitly consider the question of practical identifiability [35]. Furthermore, MCMC can be misleading in the presence of true non-identifiability [42], as can other methods such as the Bootstrap or Fisher-information-based approaches [16]. Even assuming that MCMC can reliably determine non-identifiability, MCMC can be computationally expensive, especially for models with large |*θ*|. Thus we also assess identifiability using an optimisation-based, profile likelihood approach. In contrast to previous work on temporal data and temporal processes, here we apply this approach to spatiotemporal reaction–diffusion models and spatiotemporal data. Algorithms to are available on GitHub.

A key feature of our study is that we consider two slightly different modelling scenarios. In the first scenario we assume *D* = *D*_*r*_ = *D*_*g*_ so that the diffusivity of cells in G1 phase is identical to the diffusivity of cells in the S/G2/M phase. In this case there are three unknown parameters, *θ* = (*D, k*_*r*_, *k*_*g*_). In contrast, for the second scenario we do not invoke this assumption, and there are four unknown parameters, *θ* = (*D*_*r*_, *D*_*g*_, *k*_*r*_, *k*_*g*_). Despite the relative abundance of experimental data and the apparent simplicity of the reaction-diffusion model, we find that the parameters are identifiable in the first scenario but are practically non-identifiable in the second.

Our results show that the profile likelihood provides similar results to the Bayesian MCMC approach, with the advantage of being an order of magnitude faster to implement. Typical MCMC results in Figures 2–3 require approximately 30 minutes of computation time whereas the profile likelihood results in Figure 4 require approximately one minute to compute on a standard laptop (DELL, Intel Core i7 Processor, 2.8GHz, 16GB RAM). Both methods indicate practical non-identifiability of the diffusivities in the second scenario. As mentioned, however, profile likelihood has previously been shown to be more reliable in the presence of true non-identifiability [16, 42]; this is an important consideration as the reaction–diffusion models we used here are already relatively simple and neglect certain mechanisms. Potential mechanisms that might be incorporated in further extended model include cell-cell adhesion [7, 50] or directed motion such as chemotaxis [51, 52].

Visual inspection of the posterior predictive check in Figure 2 shows that while the overall comparison between the model and the experimental data is reasonable, there are some minor discrepancies, such as in the G1 (red) subpopulation near the centre of the scratch at *t* = 32 h, suggesting some model inadequacy. This observation is consistent with the fact that we neglect some mechanisms that could potentiaily improve the quality of the match. Had we taken a more standard approach without considering an identifiability analysis, we might have been tempted to extend Equations (3)–(4) to improve the quality of match between the data and the solution of the model. Such extensions (e.g. adhesion, chemotaxis), while conceptually straightforward to implement, involve increasing the dimension of the parameter space. We caution against naively implementing such extensions since our analysis shows that even the simpler models we consider are not always practically identifiable even with the extensive data set that we work with here. A profile likelihood analysis of particular, lower-dimensional *interest parameters* within a more complex model may still be feasible, however.

There are many ways our analysis could be extended, in terms of the process model, the observation model and the experimental model. A key assumption here is that the observation model is Gaussian with a fixed variance. While this is a standard assumption [8, 13], one possible extension would be to treat the observation model parameter *σ* as a variable to be estimated at the same time as the process model parameters *θ*. Another extension would be to recognise that since we are dealing with count data, a multinomial observation model could be more appropriate. Both of these extensions could be dealt with directly in the same MCMC framework but the profile likelihood optimisation problems would become slightly more complicated than simple least squares. In terms of the process model, a further extension would be to apply the same reaction–diffusion models in a different geometry. For example, data reported by Jin et al. [53] use a different experimental approach to produce square and triangular–shaped scratches. While the same reaction–diffusion models to those used here could be used to model Jin et al.’s [53] experiments, such models would need to be solved in two spatial dimensions because of the difference in initial condition [54]. This extension could be handled by the same MCMC and the same profile likelihood approaches without increasing the dimension of the parameter space. However, this extension would increase the computational effort required to solve the reaction–diffusion model and this would amplify the computational advantage of the profile likelihood method over the Bayesian MCMC approach. In terms of the experimental model, in this work we use cell cycle labelling with two colours, red and green. More advanced cell cycle labelling technologies, such as FUCCI4 [55, 56], are now available, and these technologies label the cell cycle with four fluorescent colours [55, 56]. Data from a scratch assay where cells are labelled with FUCCI4 could be interpreted with an extended system of reaction–diffusion equations, similar to Equations (3)–(4). Such an extended system of equations would be similar to Equations (3)–(4), but the dimensionality of the parameter space would increase. With four colours we would have the possibility of working with four distinct cell diffusivities and four distinct rate parameters, thereby doubling the size of the parameter space, hence we leave this for future consideration.

## Acknowledgements

MJS is supported by the Australian Research Council (DP170100474) and the University of Canterbury Erskine Fellowship. OJM is supported by the University of Auckland, Faculty of Engineering James and Hazel D. Lord Emerging Faculty Fellowship. REB is supported by the Royal Society Wolfson Research Merit Award, the Leverhulme Trust Research Fellowship and the BBSRC (BB/R000816/1). We thank three referees for helpful feedback.

