## Supplementary Material for "Practical parameter identifiability for spatiotemporal models of cell invasion"

### 1. Numerical methods

Solutions of Equations (3)–(4) are obtained numerically. To obtain these solutions we uniformly discretise the spatial domain, $0 \leq x \leq L$, using a mesh of constant spacing, $\delta x$. Central differences are used to approximate the spatial derivatives in Equation (3)–(4), and no flux conditions at $x = 0$ and $x = L$ are approximated with a standard forward and backward difference approximation, respectively. This spatial discretisation leads to a system of $M = L/\delta x + 1$ coupled nonlinear ordinary differential equations for $r(x_i, t)$, for $i = 1, 2, \ldots, M$ and a system of $M$ coupled nonlinear ordinary differential equations for $g(x_i, t)$, for $i = 1, 2, \ldots, M$. Temporal derivatives are approximated using a backward Euler approximation with constant time step of duration $\delta t$. The resulting systems of nonlinear algebraic equations are linearised using Picard iteration and are solved sequentially, first for $r(x_i, t_j)$ and then for $g(x_i, t_j)$, using the Thomas algorithm for tridiagonal linear systems. Picard iteration is continued until the updated values for $r(x_i, t_j)$ and then $g(x_i, t_j)$ converge to within an absolute tolerance $\varepsilon$. All results correspond to $L = 1242$ $\mu$m, $\delta x = 1.0$ $\mu$m, $\delta t = 1.0$ h and $\varepsilon = 1 \times 10^{-10}$.

All numerical solutions require the specification of an initial condition, $r(x, 0)$ and $g(x, 0)$. For the results in the main document we specify these initial conditions by taking $y_i^{\mathrm{o}}$ at $t = 0$ and we use linear interpolation to determine $r(x_i, 0)$ and $g(x_i, 0)$ at each point on the finite difference mesh, $i = 1, 2, \ldots, M$.

### 2. Likelihood profiling

Here we give more details on how likelihood profiling is computationally implemented. We first give an overview of the basic concepts and show how, in our case, this methodology gives rise to a series of nonlinear least squares problems that can be solved using standard software. We then present the details of an example profiling calculation in the context of a linear regression model.

#### *2.1. Reduction to nonlinear least squares for normally-distributed observation errors*

Here we show how the profiling problem given in the main text can be reduced to solving a series of nonlinear least squares problems. We denote by $y(\psi, \lambda)$ the vector containing all model solutions $y(s_i, \psi, \lambda)$ at each observation point $s_i, i = 1, ..n$, for the parameter partition $\theta = (\psi, \lambda)$, i.e.

$$y(\psi, \lambda) = (r(s_1, \psi, \lambda), g(s_1, \psi, \lambda), ..., r(s_n, \psi, \lambda), g(s_n, \psi, \lambda)))\,. \tag{S1}$$

We assume $y^{\mathrm{o}}$ is arranged similarly. Given that the noise is iid with a normal distribution and standard deviation $\sigma$ (and is the same for observations of both red and green cells) we obtain the probability density

$$p(y^{\mathrm{o}}; \psi, \lambda) = (2\pi\sigma^2)^{-\frac{n}{2}} \exp\left(-\frac{1}{2\sigma^2} \left\|(y^{\mathrm{o}} - y(\psi, \lambda))\right\|^2\right). \tag{S2}$$

Hence, for fixed standard deviation $\sigma$, the profile (normalised) likelihood for $\psi$ is

$$\mathcal{L}_p(\psi; y^{\mathrm{o}}) = \frac{\sup_\lambda \left(\exp\left(-\frac{1}{2\sigma^2} \|(y^{\mathrm{o}} - y(\psi, \lambda))\|^2\right)\right)}{\sup_{(\psi,\lambda)} \left(\exp\left(-\frac{1}{2\sigma^2} \|(y^{\mathrm{o}} - y(\psi, \lambda))\|^2\right)\right)}, \tag{S3}$$

where the common factor $(2\pi\sigma^2)^{-\frac{n}{2}}$ cancels. As maximisation and taking the exponential commute, we have

$$\mathcal{L}_p(\psi; y^{\mathrm{o}}) = \frac{\exp\left(\sup_\lambda \left(-\frac{1}{2\sigma^2} \|(y^{\mathrm{o}} - y(\psi, \lambda))\|^2\right)\right)}{\exp\left(\sup_{(\psi,\lambda)} \left(-\frac{1}{2\sigma^2} \|(y^{\mathrm{o}} - y(\psi, \lambda))\|^2\right)\right)}, \tag{S4}$$

and hence the key computations involve solving

$$\sup_\lambda -\frac{1}{2\sigma^2} \left\|(y^{\mathrm{o}} - y(\psi, \lambda))\right\|^2 = \inf_\lambda \frac{1}{2\sigma^2} \left\|(y^{\mathrm{o}} - y(\psi, \lambda))\right\|^2 \tag{S5}$$

for each value of $\psi$. Equation (S5) is a nonlinear least squares problem that can be solved using standard numerical software. In this work we choose to solve such problems using the lsqnonlin routine in MATLAB [1]. After solving each least squares problem we take exponentials to regain the appropriate likelihoods. The overall optimisation for the denominator can be computed by a subsequent optimisation over the fixed $\psi$ sub-problems, as $\sup_{(\psi,\lambda)} f(\psi,\lambda) = \sup_{\psi}\left(\sup_{\lambda}\left(f(\psi,\lambda)\right)\right)$ for any function $f$.

#### 2.2. *Profiling example: linear regression*

Here we consider profiling for the simple linear (in parameters) model

$$y_i = f(x_i, a, b) + \epsilon_i = ax_i + bx_i^2 + \epsilon_i, \quad \epsilon_i \sim \mathcal{N}(0, 1), \tag{S6}$$

i.e. $y_i \sim \mathcal{N}(f(x_i, a, b), 1)$, for $i = 1, ...N$, meaning that we have Gaussian observation model with $\sigma = 1$. We wish to compute the profiles $\mathcal{L}_p(a; y^{\rm o})$ and $\mathcal{L}_p(b; y^{\rm o})$, where $y^{\rm o}$ denotes the combined vector of observations.

Following the arguments outlined above, the key computations for $\mathcal{L}_p(a; y^{\rm o})$, for example, involve solving least squares problems of the form

$$\inf_{b} \frac{1}{2\sigma^2} \left\|(y^{\rm o} - f(a, b))\right\|^2 \tag{S7}$$

for a grid of $a$ values, and where model outputs at all $x$ observation locations have been collected (as before) into a vector, here denoted by $f(a, b)$. The relative likelihood function is then computed by taking the maximum across all $a$ values to obtain the overall minimiser for the denominator, and then taking exponentials to regain the likelihoods. We assume that the true likelihood is sufficiently smooth so that we can extend the likelihood as calculated at gridded values to a likelihood defined at all values via interpolation between grid points. Similarly, we compute $\mathcal{L}_p(b; y^{\rm o})$ by switching the roles of $a$ and $b$ as interest and nuisance parameters.

A simple example of profiling the two-dimensional likelihood function corresponding to the model in Equation (S6) is shown Figure S1. Here we generated 40 samples from the model over a regularly spaced grid for $x \in [0, 1]$. The true parameters used were $a = 1, b = 3$. The two-dimensional likelihood was generated by direct calculation over a $200 \times 200$ grid.

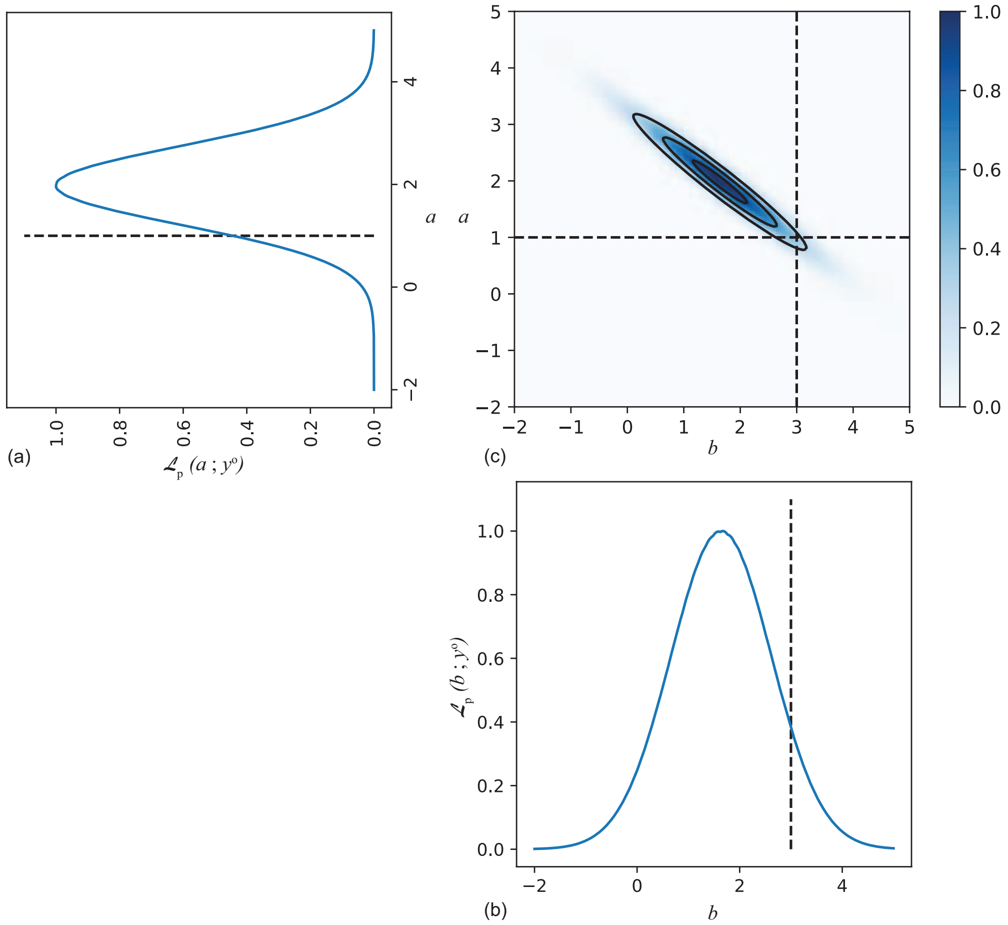


Figure S1: Example of profiling the two-dimensional likelihood function corresponding to the model S6 to obtain profile likelihoods for each parameter $a$ (a) and $b$ (b). Here we generated 40 samples from the model over a regularly spaced grid for $x \in [0, 1]$. The true parameters used were $a = 1, b = 3$ and these are indicated by dashed lines. The observation model is Gaussian with $\sigma = 1$. Each profile corresponds to the maximum taken over the other dimension and hence to a 'profile' view of the two-dimensional likelihood (c). The two-dimensional likelihood is evaluated over a $200 \times 200$ grid.

### 3. Additional MCMC results for the experimental data

Results in Figures 2-3 report the Bayesian MCMC results for the two scenarios where $D = D_r = D_g$ and $D_r \neq D_g$, respectively. In those results we present a visual posterior predictive check. Additional results in Figure S2–S3 show identical figures except that instead of the posterior predictive check we compare the experimental data with the solution of the model using the univariate mode for the case where $D = D_r = D_g$ (Figure S2) and the median for the case where $D_r \neq D_g$ (Figure S3)

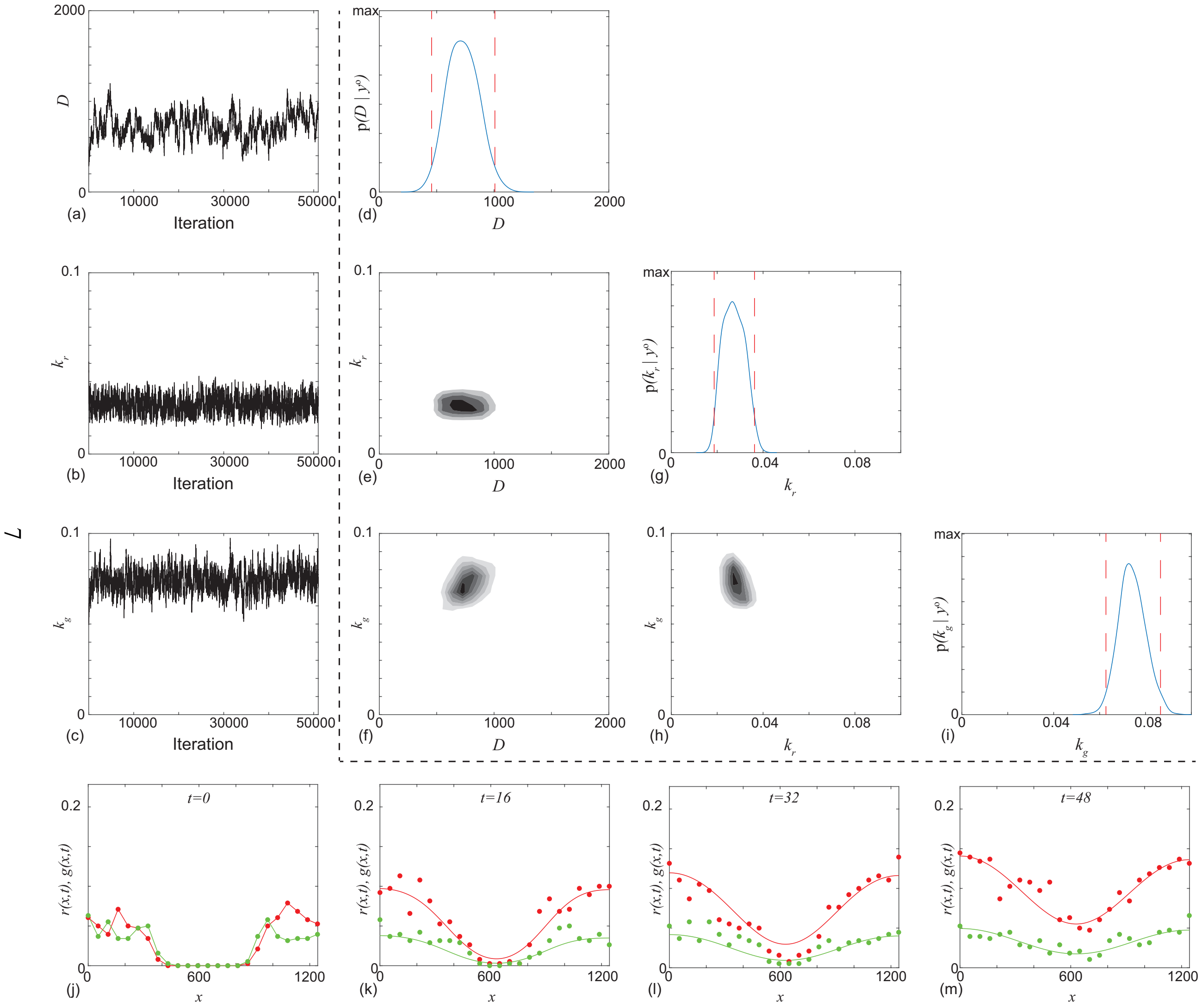


Figure S2: Typical Markov chain iterations, of length 51,000, for $D$, $k_r$ and $k_g$ in (a)–(c), respectively. In this case the Markov chain is initiated with $\theta_0 = (500, 0.05, 0.05)$. Results in (d)–(i) show a plot matrix representation of the univariate marginals and bivariate marginals estimated using the final 50,000 iterations of the Markov chain in (a)–(c). For the univariate distribution the posterior modes are $\bar{D} = 707\ \mu\text{m}^2$ /h, $\bar{k}_r = 0.0266$ /h, and $\bar{k}_g = 0.0727$ /h, and the 95% credible intervals are $D \in [455, 1007]$, $k_r \in [0.0187, 0.0362]$ and $k_g \in [0.0628, 0.0727]$. In the univariate marginals the 95% credible intervals are shown in red dashed vertical lines, in the bivariate marginals the region of maximum density is shown in the darkest shade. Results in (j)–(m) superimpose the solution of Equations (1)–(2) with the median $\theta = (707, 0.0266, 0.0727)$ superimposed on the experimental data. MCMC results use $\sigma = 0.05$.

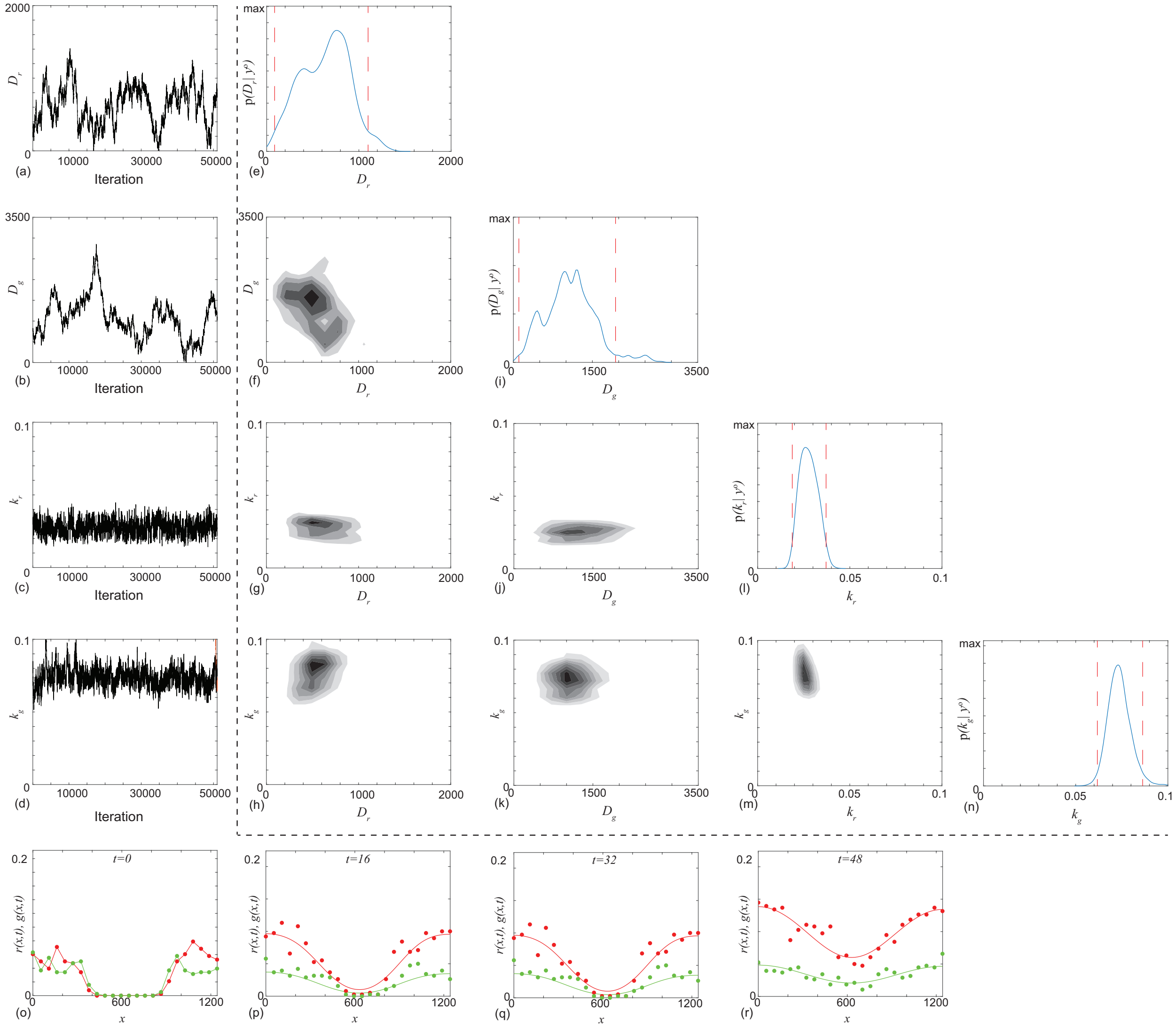


Figure S3: Typical Markov chain iterations, of length 51,000, for $D_r$, $D_g$, $k_r$ and $k_g$ in (a)–(d), respectively. In this case the Markov chain is initiated with $\theta_0 = (228, 803, 0.0334, 0.0790)$. Results in (e)–(n) show a plot matrix representation of the univariate marginals and bivariate marginals estimated using the final 50,000 iterations of the Markov chain in (a)–(d). For the univariate distribution the posterior modes are $\bar{D}_r = 758$ $\mu$m$^2$ /h, $\bar{D}_g = 1203$ $\mu$m$^2$ /h, $\bar{k}_r = 0.0259$ /h, and $\bar{k}_g = 0.0703$ /h. The 95% credible intervals are $D_r \in [86, 1100]$, $D_g \in [104, 1939]$, $k_r \in [0.0188, 0.0371]$ and $k_g \in [0.0617, 0.0863]$. In the univariate marginals the 95% credible intervals are shown in red dashed vertical lines, in the bivariate marginals the region of maximum density is shown in the darkest shade. Results in (o)–(r) superimpose the solution of Equations (1)–(2) with the median $\theta = (660, 1063, 0.0274, 0.0733)$ superimposed on the experimental data. MCMC results use $\sigma = 0.05$.

As noted in the main document, the Markov chain in Figure 2a indicates that the first scenario with $D = D_r = D_g$ is identifiable because the trace plot appears to converge whereas results in Figure 3a-b indicate practical non-identifiability in the second scenario because the trace plots do not converge for $D_r$ and $D_g$. Since the MCMC results in Figure 2 and Figure 3 are for one particular choice of $\theta_0$, we provide additional results here to explore the impact of varying $\theta_0$. Additional results in Figures S4–S5 provides further evidence to support these conclusions by showing additional MCMC results for different choices of $\theta_0$ for the first case where $D = D_r = D_g$ (Figure S4) and in the second case where $D_r \neq D_g$ (Figure S5). Results in Figure S4 indicate rapid convergence for $D$, $k_r$ and $k_g$ regardless of $\theta_0$. In contrast, results in Figure S5 indicate rapid convergence for $k_r$ and $k_g$ regardless of $\theta_0$, but we see that the Markov chains for $D_r$ and $D_g$ depend upon $\theta_0$ and do not converge.

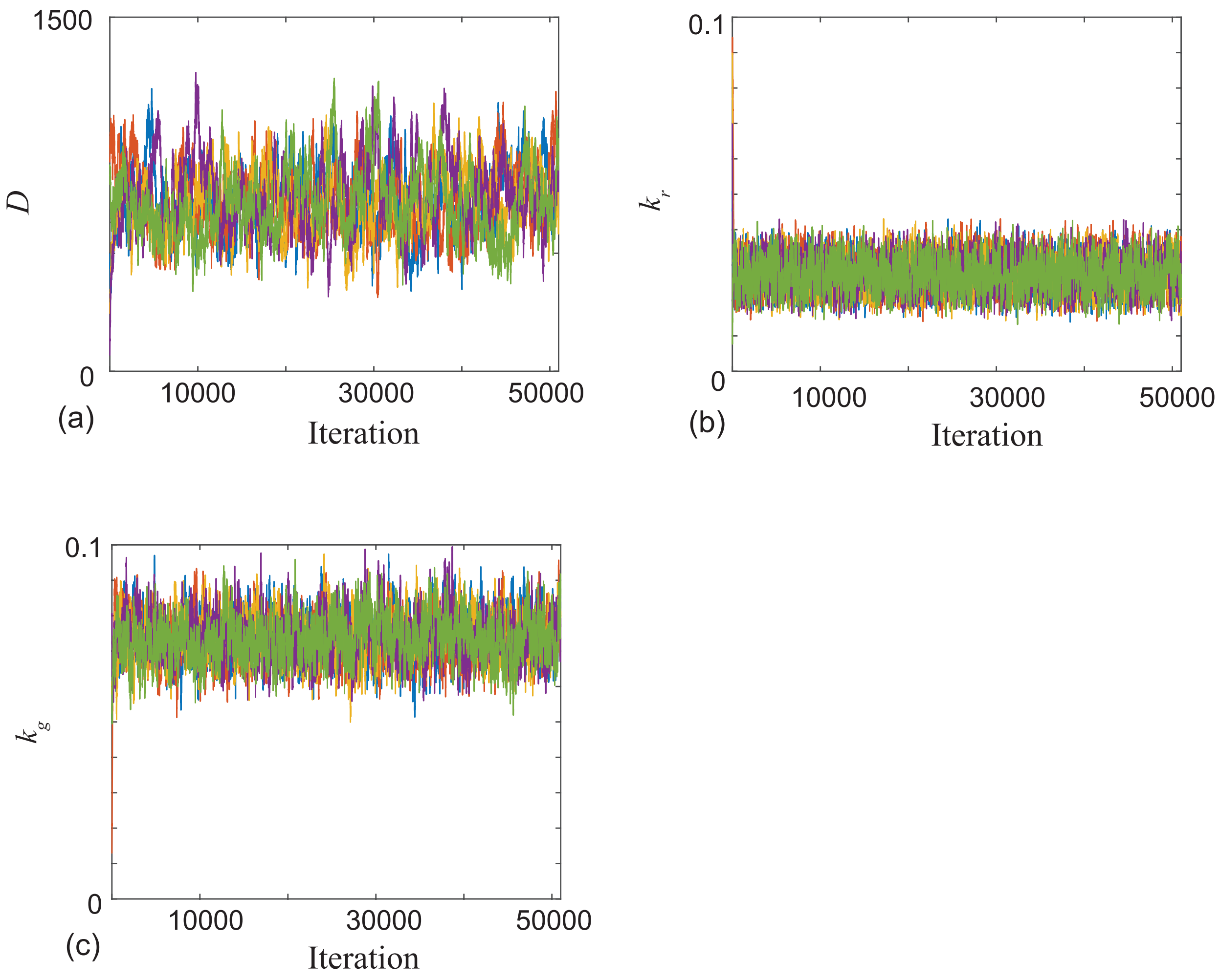


Figure S4: Five realizations of the Markov Chain from Figure 2. The initial choice of $\theta_0$ are (500, 500, 0.0500, 0.0500) (blue), (906, 0.0900, 0.0130) (orange), (259, 0.0900, 0.0600) (yellow), (67, 0.0700, 0.0800) (purple) and (866, 0.0096, 0.0550) (green). Results in Figure 2 correspond to the blue chain.

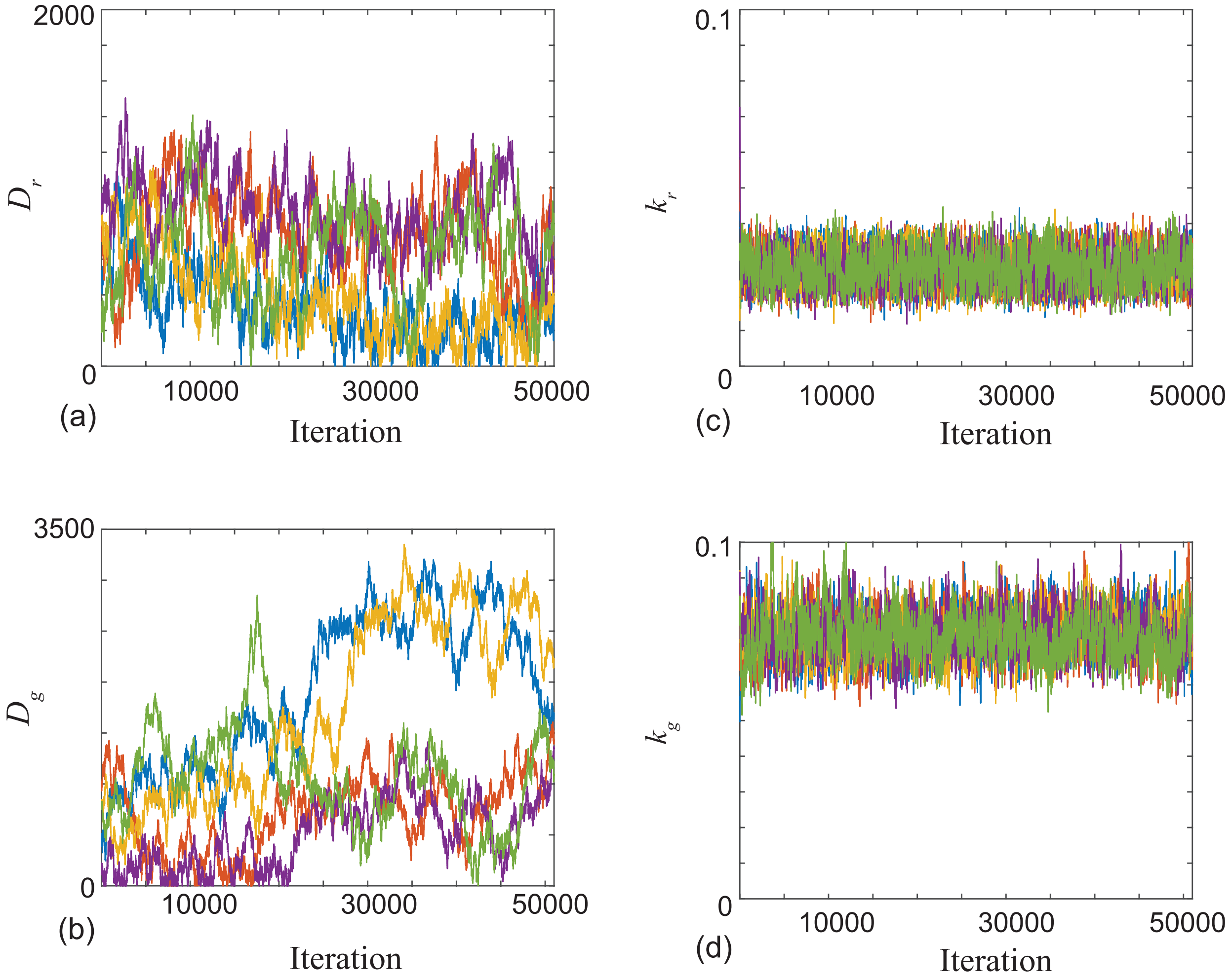


Figure S5: Five realizations of the Markov Chain from Figure 3. The initial choice of $\theta_0$ are $(500, 500, 0.0500, 0.0500)$ (blue), $(705, 979, 0.0573, 0.0875)$ (orange), $(905, 814, 0.0127, 0.0913)$ (yellow), $(905, 215, 0.0727, 0.0914)$ (purple) and $(228, 803, 0.0334, 0.0790)$ (green). Results in Figure 3 correspond to the green chain.

### 4. Additional profile likelihood results for the experimental data

Results in Figure S6 show the profile for the experimental data under the second scenario where $D_r \neq D_g$. Here the interest parameter is the difference in diffusivities, i.e. $\psi = D_r - D_g$. Results indicate that compatible results include zero, but also differences at least as large as +/− 1500, and the cut-off 0.15 lies outside of this interval.

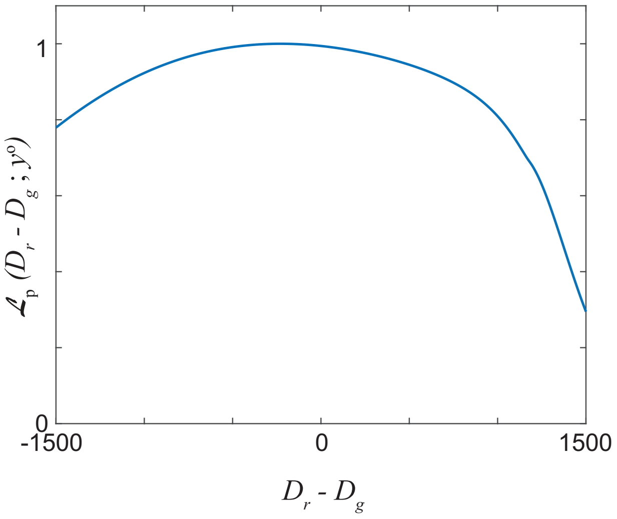


Figure S6: Profile likelihood for $D_r - D_g$ for the second scenario where $D_r \neq D_g$. The profile likelihood is computed using a grid of 50 equally-spaced points; cubic interpolation is used for display and to determine confidence intervals.

### 5. Additional MCMC results for finely resolved synthetic data

To aid in demonstrating the key difference between practical and structural identifiability we now repeat all results from the main document using synthetic data for which we specify known parameter values. This approach allows us to collect as much data as we prefer since we are not limited by experimental constraints. Since we can collect as much data as we like we will refer to these results as *finely resolved synthetic data*. This synthetic data corresponds to solutions of Equations (1)–(2) on the same domain. The initial condition is $r(x, 0) = g(x, 0) = 0.2$ for $0 < x < 400$, $r(x, 0) = g(x, 0) = 0.2$ for $842 < x < 1242$, and $r(x, 0) = g(x, 0) = 0$ for $400 < x < 842$. We consider two scenarios, the first has $D_r = D_g$ with $\theta = (500, 500, 0.05, 0.10)$, and the second has $D_r \neq D_g$ with $\theta = (700, 300, 0.05, 0.10)$. In both scenarios we collect synthetic data at each point on the finite difference mesh, and at $t = 0, 16, 32$ and 48. Thus $y^o$ consists of $N = 1243 \times 4 = 4972$ observations of both of $r$ and $g$, that is 9944 measurements of density in total. Therefore the synthetic data cases involves dealing with almost two orders of magnitude more observations than was possible with the experimental data. In the next section we also consider using a finer temporal resolution, along with the finer spatial resolution, using profile likelihood.

Results in Figures S7-S8 show the Bayesian MCMC results for the two scenarios respectively. In both cases we see that the Markov chains appear to converge and the univariate posterior distributions are regular shaped with clearly defined maxima that are close to the expected values. These results indicate that the parameters are identifiable in both synthetic scenarios provided we have access to sufficient observations, i.e. we expect that with even more data our estimates will converge to the true values. However, even in this dense data limit, when we compare results in Figure S7 for $D_r = D_g$ with results in Figure S8 for $D_r \neq D_g$, we see that the credible intervals in second scenario are relatively wider than in the first scenario. This indicates that identifiability in the second scenario is more challenging than in the first, and this is consistent with the results in Figure 2-3 for the experimental data. Nonetheless, with this finely resolved synthetic data we see that the MCMC results leads to well-formed posteriors and each marginal posterior is approximately symmetric and unimodal with the point estimates close to the known values in each case.

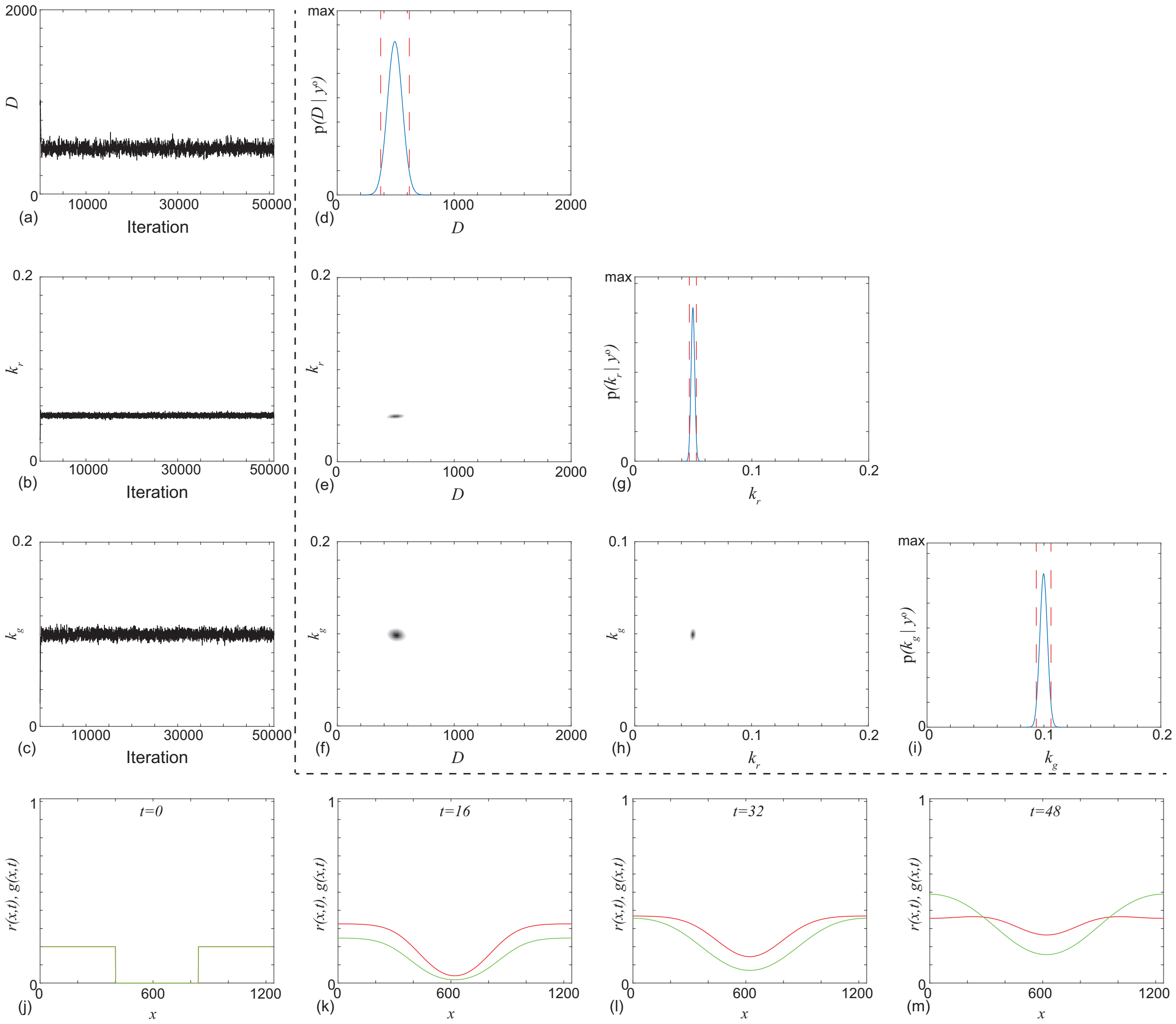


Figure S7: Typical Markov chain iterations, of length 51,000, for $D$, $k_r$ and $k_g$ in (a)–(c), respectively. In this case the Markov chain is initiated with $\theta = (1000, 0.025, 0.025)$. In this case the true value is $\theta = (500, 0.050, 0.10)$, Results in (d)–(i) show a plot matrix representation of the univariate marginals and bivariate marginals estimated using the final 50,000 iterations of the Markov chain in (a)–(c). For the univariate distribution the posterior modes are $\bar{D} = 493\ \mu\mathrm{m}^2$ /h, $\bar{k}_r = 0.0496$ /h, and $\bar{k}_g = 0.0997$ /h, and the 95% credible intervals are $D \in [372, 517]$, $k_r \in [0.0466, 0.0526]$ and $k_g \in [0.0935, 0.1061]$. In the univariate marginals the 95% credible intervals are shown in red vertical dashed lines, in the bivariate marginals the region of maximum density is shown in the darkest shade. Results in (j)–(m) show the solution of Equations (1)–(2) with $\theta = (1000, 0.025, 0.025)$. MCMC results use $\sigma = 0.5$.

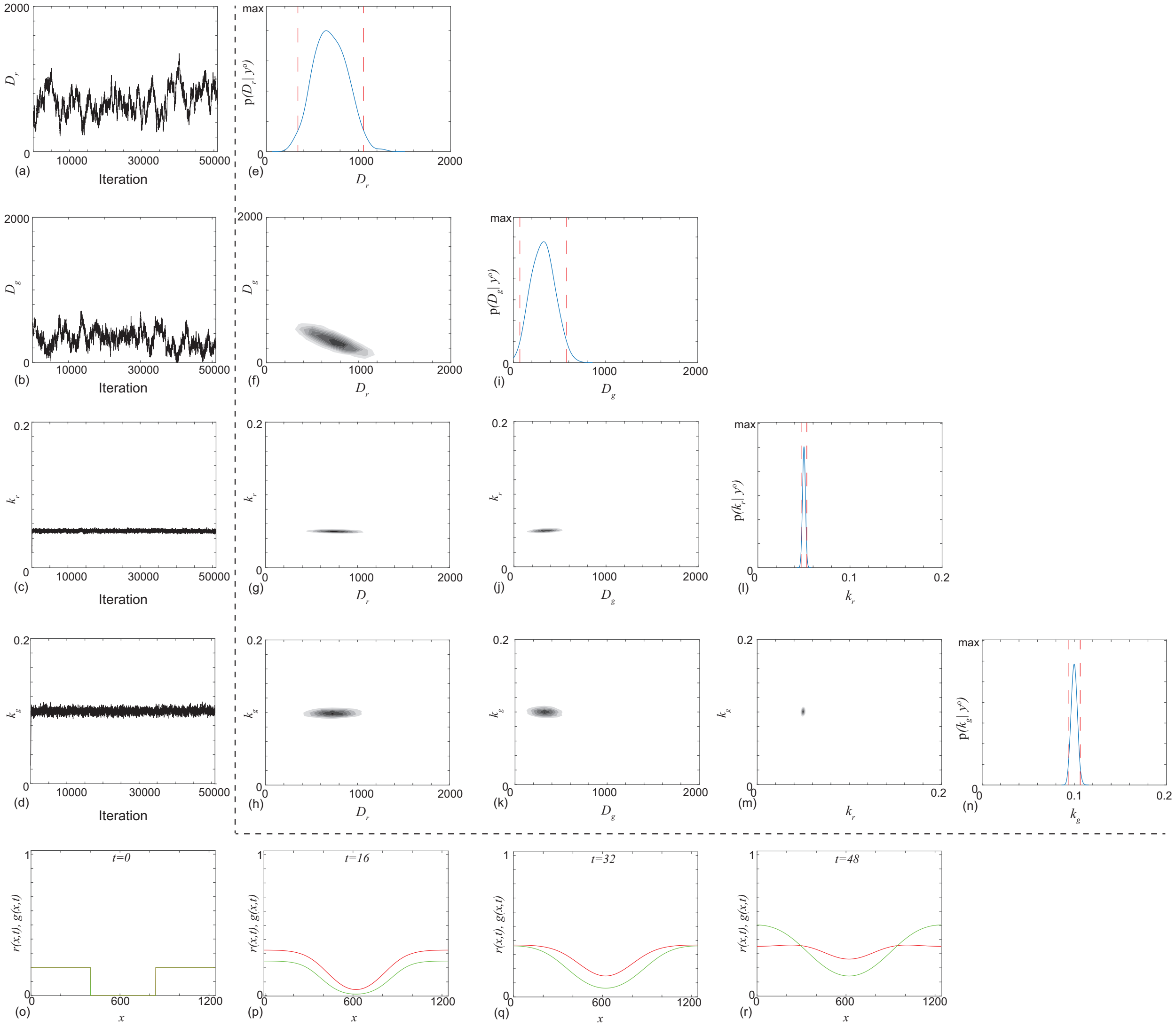


Figure S8: Typical Markov chain iterations, of length 51,000, for $D_r$, $D_g$, $k_r$ and $k_g$ in (a)–(d), respectively. In this case the Markov chain is initiated with $\theta = (500, 500, 0.0250, 0.0250)$. In this case the true value is $\theta = (700, 300, 0.050, 0.10)$ Results in (e)–(n) show a plot matrix representation of the univariate marginals and bivariate marginals estimated using the final 50,000 iterations of the Markov chain in (a)–(d). For the univariate distribution the posterior modes are $\bar{D}_r = 658\ \mu\text{m}^2$ /h, $\bar{D}_g = 320\ \mu\text{m}^2$ /h, $\bar{k}_r = 0.0501$ /h, and $\bar{k}_g = 0.0999$ /h. The 95% credible intervals are $D_r \in [342, 1056]$, $D_g \in [69, 577]$, $k_r \in [0.0470, 0.03532]$ and $k_g \in [0.0933, 0.1064]$. In the univariate marginals the 95% credible intervals are shown in red vertical dashed lines, in the bivariate marginals the region of maximum density is shown in the darkest shade. Results in (o)–(r) show the solution of Equations (1)–(2) with $\theta = (700, 300, 0.050, 0.100)$. MCMC results use $\sigma = 0.5$.

### 6. Additional profile results for synthetic data

Results in Figures S9-S10 show additional profiling results for the first and second scenarios discussed above, i.e. for $D_r = D_g$, with $\theta = (500, 500, 0.05, 0.10)$, and for $D_r \neq D_g$, with $\theta = (700, 300, 0.05, 0.10)$. As above, here we use finely resolved synthetic data, though now we also take finer measurements in time: we take observations at each spatial location of the solution grid, as well as observations at each hour between $t = 0$ and $t = 48$ h. This gives $N = 1243 \times 48 = 59664$ observations (excluding the initial condition) of both of $r$ and $g$, that is 119328 measurements of density in total. In Figure S11 we also consider data collected every hour but using the coarse experimental grid, given parameters set according to the second scenario. For all results we used the same measurement precision, $\sigma = 0.05$, as used for the experimental data.

In the first scenario where $D_r = D_g$ we see sharply peaked profiles for $D_r$, $D_g$, $k_r$ and $k_g$ centred at the known values (Figure S9a-b, d-e). We also profile for $D_r - D_g$ and obtain sharp profiles with a clear defined peak near zero, as expected (Figure S9c). In the second scenario where $D_r \neq D_g$ and $D_r - D_g = 400$, and for high resolution spatiotemporal data (Figure S10), we again see sharply peaked profiles centred at the known values. The maximum likelihood estimates correspond to the true values in each scenario, i.e. $\theta^* = (500, 0.050, 0.100)$ and $\theta^* = (700, 300, 0.050, 0.100)$, respectively.

In the case of high temporal resolution but coarse spatial resolution, we see better results than those from the experimental data, but that it is still difficult to distinguish $D_r$ and $D_g$ (Figure S11). For example, the approximate 95% confidence interval for the difference $D_r - D_g$ is [-436, 902], whereas the real difference is 400.

These numerical results indicate that the model parameters are likely structurally identifiable, i.e. that the estimates can be expected to converge to the true values in the limit of increasing amounts of spatiotemporal data. The convergence rates of the diffusion coefficients, and the convergence rates of the difference in diffusion coefficients, appear to be slower than those for the rate parameters, however, thus indicating that they are more difficult to estimate in general. This is consistent with the practical identifiability results with real data in the main text. We also see that, while the finest resolution results indicate that the model is structurally identifiable, high temporal resolution by itself does not appear to be sufficient for good practical identifiability.

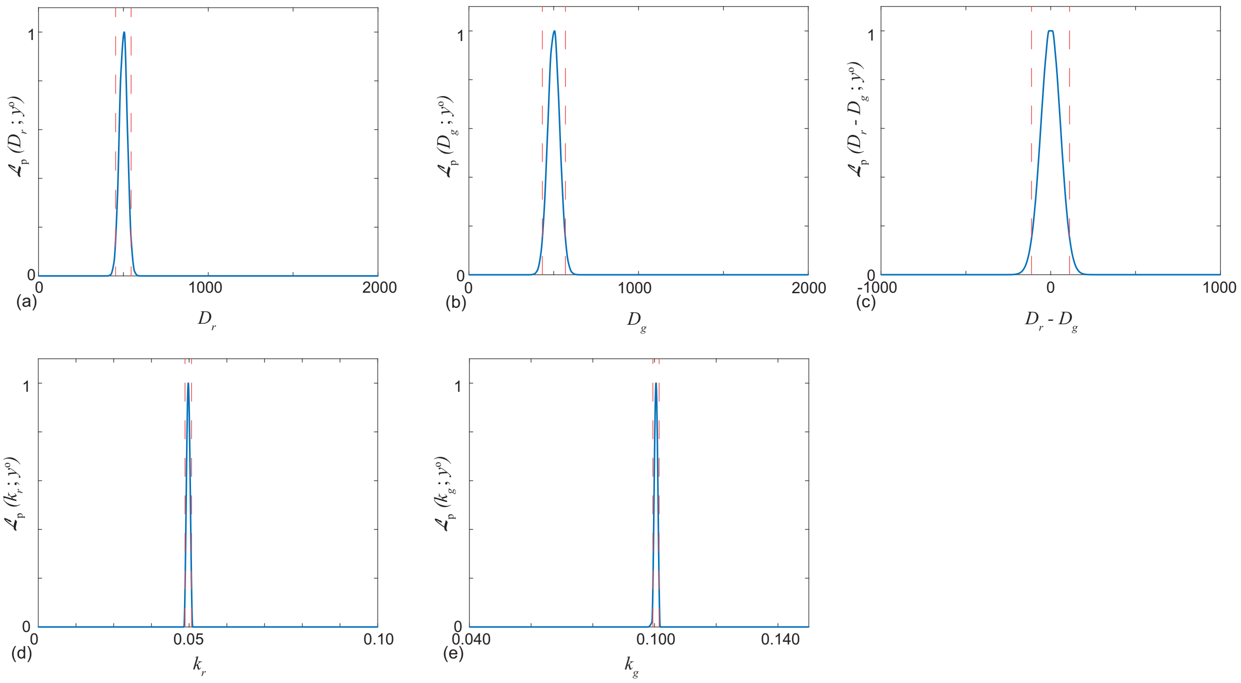


Figure S9: Profile likelihoods for synthetic data with $\theta = (500, 500, 0.05, 0.10)$, representing increasing amounts of spatiotemporal data. Measurements are collected at each spatial and temporal point of the numerical solution, between $t = 0$ to $t = 48$ h. Each profile likelihood is computed using a grid of 100 equally-spaced points for the target parameter; cubic interpolation is used for display and to determine confidence intervals. Approximate 95% confidence intervals are indicated based on a relative likelihood cutoff of 0.15 [2]. Interval estimates are: $D_r \in$ [454, 545], $D_g \in$ [433, 569], $D_r - D_g \in$ [-113, 111], $k_r \in$ [0.0489, 0.0506] and $k_g \in$ [0.0995, 0.1015]. Maximum likelihood estimates are: $D_r^*$, $D_g^*$, $(D_r - D_g)^*$, $k_r^*$, $k_g^*$ = 500, 500, 0.0, 0.05, 0.10. Our estimates can thus be seen to be converging to the true values in the limit of increasing amounts of spatiotemporal data. Results used $\sigma = 0.05$.

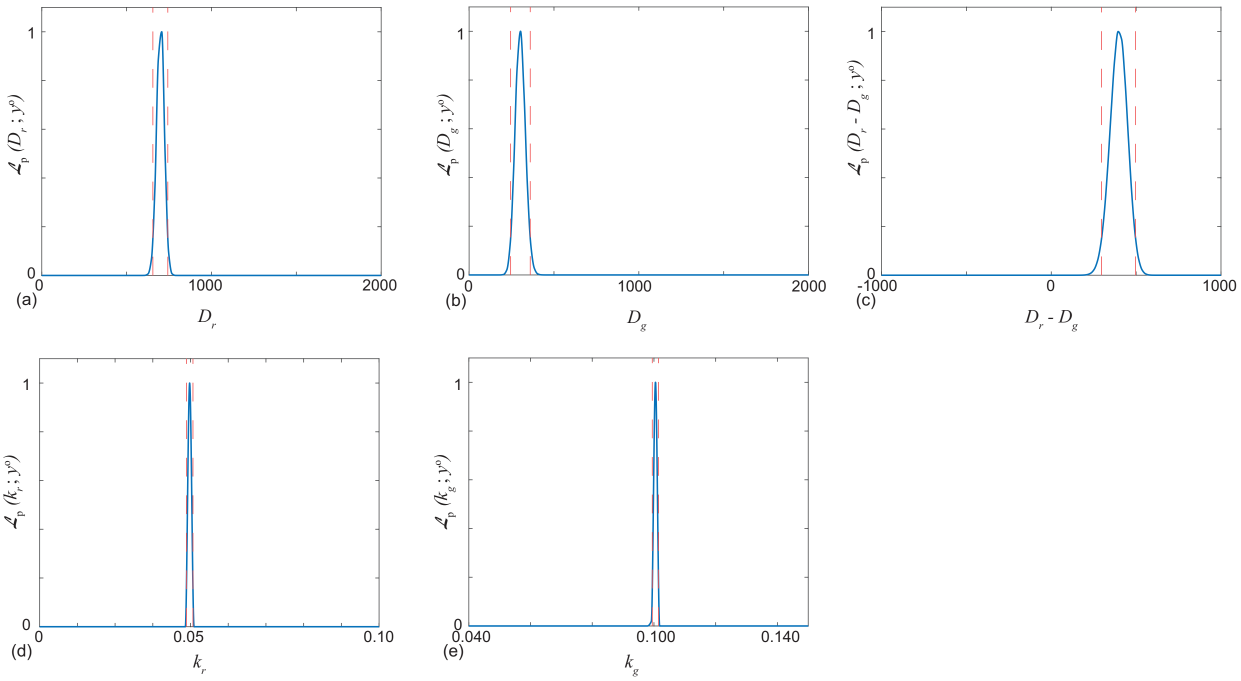


Figure S10: Profile likelihoods for synthetic data with $\theta = (700, 300, 0.05, 0.10)$ , representing increasing amounts of spatiotemporal data. Measurements are collected at each spatial and temporal point of the numerical solution, between $t = 0$ to $t = 48$ h. Each profile likelihood is computed using a grid of 100 equally-spaced points; cubic interpolation is used for display and to determine confidence intervals. Approximate 95% confidence intervals are indicated based on a relative likelihood cutoff of 0.15 [2]. Interval estimates are: $D_r \in [655, 743]$, $D_g \in [245, 360]$, $D_r - D_g \in [296, 497]$, $k_r \in [0.0489, 0.0506]$and $k_g \in [0.0995, 0.1015]$. Maximum likelihood estimates are: $D_r^*$, $D_g^*$, $(D_r - D_g)^*$, $k_r^*$, $k_g^*$ = 700, 300, 400, 0.05, 0.10. Our estimates can thus be seen to be converging to the true values in the limit of increasing amounts of spatiotemporal data. Results used $\sigma = 0.05$.

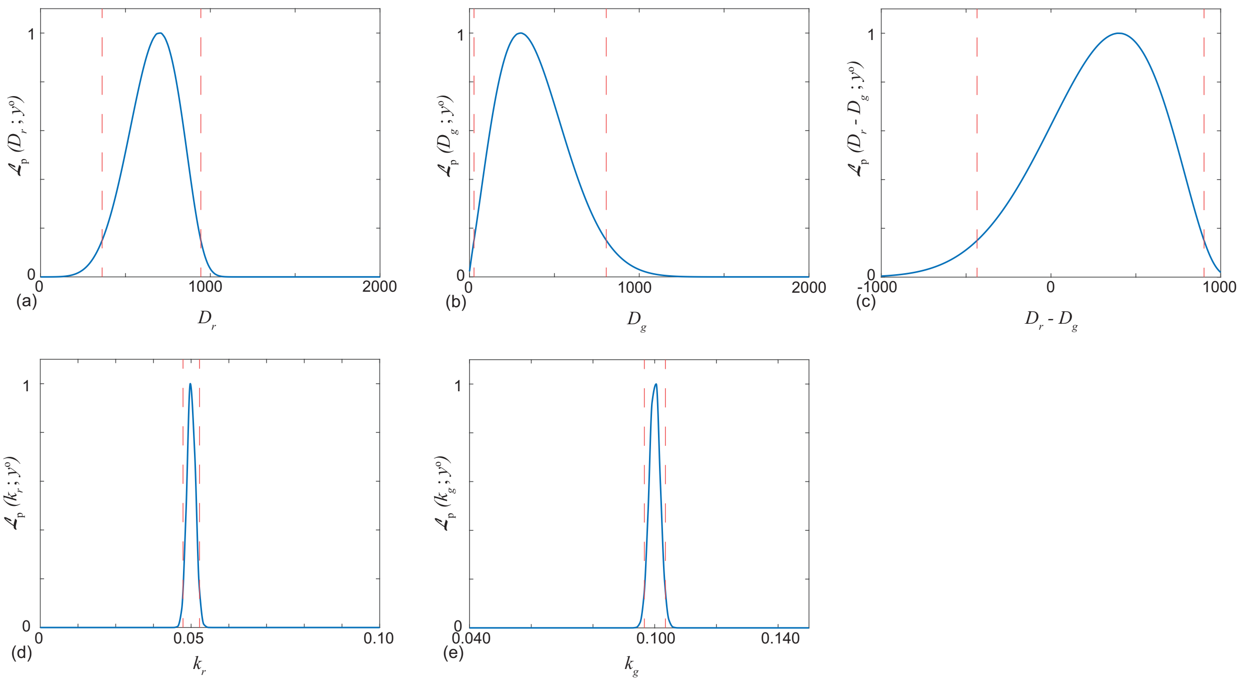


Figure S11: Profile likelihoods for synthetic data with $\theta = (700, 300, 0.05, 0.10)$, representing the case of increasing temporal resolution and fixed, experimental spatial resolution. Measurements are collected every hour between $t = 0$ to $t = 48$ at the same spatial grid points as the experimental results in the main text. Each profile likelihood is computed using a grid of 100 equally-spaced points; cubic interpolation is used for display and to determine confidence intervals. Approximate 95% confidence intervals are indicated based on a relative likelihood cutoff of 0.15 [2]. Interval estimates are: $D_r \in$ [363, 944], $D_g \in$ [26.4, 805], $D_r - D_g \in$ [-436, 902], $k_r \in$ [0.0478, 0.0522] and $k_g \in$ [0.0966, 0.1035]. Maximum likelihood estimates are: $D_r^*$, $D_g^*$, $(D_r - D_g)^*$, $k_r^*$, $k_g^*$ = 700, 300, 400, 0.05, 0.10. Our estimates can thus be seen to be converging to the true values in the limit of increasing amounts of temporal data, but at a slower rate than when spatial resolution is also increased. Results used $\sigma = 0.05$.

[1] Mathworks. 2020. lsqnonlin. Solve nonlinear least-squares. https://au.mathworks.com/help/optim/ug/lsqnonlin.html. (Accessed January 2020).

[2] Pawitan Y. 2001. In all likelihood: statistical modelling and inference using likelihood. Oxford University Press.
